# UBR5 sustains effector regulatory T cell fitness by restraining the p53 checkpoint

**DOI:** 10.64898/2026.11.25.769189

**Authors:** Aizhang Xu, Chenyi Yang, Emily Xie, Haijun Zhang, Tuo Zhang, Shannon M Buckley, Gang Lin, Pengbo Zhou, Roberta Zappasodi, Xiaojing Ma

## Abstract

Regulatory T cells (Tregs) restrain antitumor immunity, yet the mechanisms that sustain their effector functions remain incompletely understood. Here, we identify the E3 ubiquitin ligase UBR5 as a key regulator of effector Treg (eTreg) fitness. Conditional and mosaic genetic models reveal that UBR5 supports Treg proliferation, effector differentiation, and suppressive function. Treg-specific UBR5 deletion inhibits tumor growth and sensitizes tumors to PD-1 blockade. Mechanistically, UBR5 promotes p53 protein turnover through its HECT domain, limiting activation of the p53–p21 checkpoint. Genetic deletion of p53 or p21 rescues key defects in UBR5-deficient Tregs. Single-cell transcriptomic profiling reveals impaired effector-associated transcriptional programs in UBR5-deficient tumor Tregs. Analysis of human tumors further reveals an association of UBR5 expression with eTreg features and nonresponse to immune checkpoint blockade. Together, these findings identify a mechanism linking ubiquitin-dependent protein homeostasis to Treg proliferative fitness and immune suppression, with implications for cancer immunotherapy.

**Summary:** Xu et al. identify the E3 ubiquitin ligase UBR5 as a key regulator of effector regulatory T cell fitness. UBR5 restrains the p53–p21 checkpoint to support Treg proliferation and immune suppression. Its deletion enhances antitumor immunity, while human tumor data associate UBR5 expression with effector Treg features and immunotherapy nonresponse.

## Introduction

Regulatory T cells (Tregs) are essential for maintaining immune homeostasis (Sakaguchi et al., 2008), but in cancer they represent a major barrier to effective antitumor immunity (Tay et al., 2023). Quiescent resting Treg (rTreg) cells reside in lymphoid tissues to prevent lymphoproliferation (Bergot et al., 2015; Luo et al., 2016). Upon antigen stimulation, rTreg cells enter cell cycle and upregulate the expression of Treg effector molecules (e.g. IL-10, LAG3, CD73, GITR, ICOS) and specific integrins and chemokine receptors (e.g. CD103, CCR6, CCR8) (Munoz-Rojas and Mathis, 2021). rTreg cells eventually differentiate into effector (eTreg) cells which migrate to peripheral tissues and inhibit excess inflammatory response to maintain tissue homeostasis. Coordinated regulation of Treg proliferation and effector differentiation is therefore critical for maintaining the functional Treg compartment and peripheral immune tolerance. In the context of tumor progression, eTregs infiltrate tumors to suppress antitumor immunity (Sakaguchi et al., 2020). Tumor-infiltrating Tregs accumulate at high frequency and adopt a highly activated, effector-like state characterized by sustained proliferation, metabolic adaptation, and potent suppressive function (Swatler et al., 2024). This tumor-adapted Treg program enables durable immune suppression even in the face of chronic antigen stimulation and inflammatory stress, and it is strongly associated with resistance to immune checkpoint blockade across multiple cancer types. Tumor-infiltrating Tregs can adopt a fragile state in which they retain Foxp3 expression but lose suppressive capacity and acquire inflammatory features such as IFN-γ production. This state, driven by tumor-associated inflammatory cues, enhances antitumor immunity without systemic autoimmunity and represents a distinct regulatory checkpoint from Treg depletion or lineage instability (Overacre-Delgoffe et al., 2017; Zhou et al., 2009). Although prior work has identified transcriptional and epigenetic modules governing eTreg differentiation (Alvisi et al., 2020; Chowdhary et al., 2024), the mechanisms that couple proteostasis to Treg cell cycle, proliferation and stability remain incompletely defined. This gap limits the development of strategies that selectively destabilize intratumoral Tregs without compromising systemic tolerance.

Ubiquitin-mediated proteostasis plays a central role in coordinating cell-cycle control, stress responses, and lineage stability across immune cell types. Among E3 ubiquitin ligases, UBR5 (ubiquitin protein ligase E3 component N-recognin 5) has been implicated in diverse cellular processes, including DNA damage responses (Gudjonsson et al., 2012), transcriptional regulation (Mark et al., 2023), and cell-cycle progression (Smits, 2012; Zhang et al., 2023b), and is frequently dysregulated in cancer. While tumor-intrinsic roles for UBR5 in promoting growth and immune evasion have been described (Liao et al., 2017; Shearer et al., 2015; Song et al., 2020a; Song et al., 2020b), whether UBR5 functions within immune cells—and in regulatory T cells in particular—has remained largely unexplored. Given the exceptional proliferative demands and stress tolerance required of eTregs, we hypothesized that UBR5 may contribute to the molecular mechanisms that sustain the effector program and function of regulatory T cells.

To define the role of UBR5 in regulatory T cells in vivo, we employed conditional and mosaic genetic approaches to selectively disrupt UBR5 function within the Treg lineage. These strategies allowed us to assess the cell-intrinsic requirement for UBR5 during Treg proliferation and effector differentiation under both steady-state and tumor-associated conditions, while minimizing confounding effects from developmental defects or systemic immune perturbation. By integrating these genetic models with transcriptomic and single-cell analyses, as well as functional tumor assays, we systematically examined how UBR5 shapes eTreg fitness and suppressive programs within the tumor microenvironment (TME).

## Results

### UBR5 intrinsically supports effector Treg differentiation and suppressive function

To determine whether UBR5 might influence Treg biology, we first examined its expression across human immune tissues (Kim et al., 2014) and found that UBR5 protein is highly expressed in both B and T lymphocytes (Figure S1A). Analysis of mouse RNA-seq datasets showed that *Ubr5* is enriched in peripheral CD8^+^ intraepithelial lymphocytes (IEL) and colonic Foxp3^+^ Tregs (Figure S1B). These findings suggested a potential role for UBR5 in Tregs. We therefore crossed *Ubr5^f/f^* mice with *Foxp3^YFP-Cre^* mice to restrict UBR5 deletion to the Treg lineage (UBR5 cKO). Western blot analysis confirmed efficient UBR5 deletion in Tregs and preserved expression in conventional T (Tconv) cells (Figure 1A). Interestingly, UBR5 protein expression was higher in wild-type Tregs than in Tconv cells, suggesting a Treg-enriched role. Furthermore, Tconv cells from UBR5 cKO mice exhibited elevated UBR5 protein compared with those from WT mice, suggesting possible compensatory changes in UBR5 expression or altered homeostasis in the setting of Treg dysfunction due to UBR5 loss.

**Figure 1.**
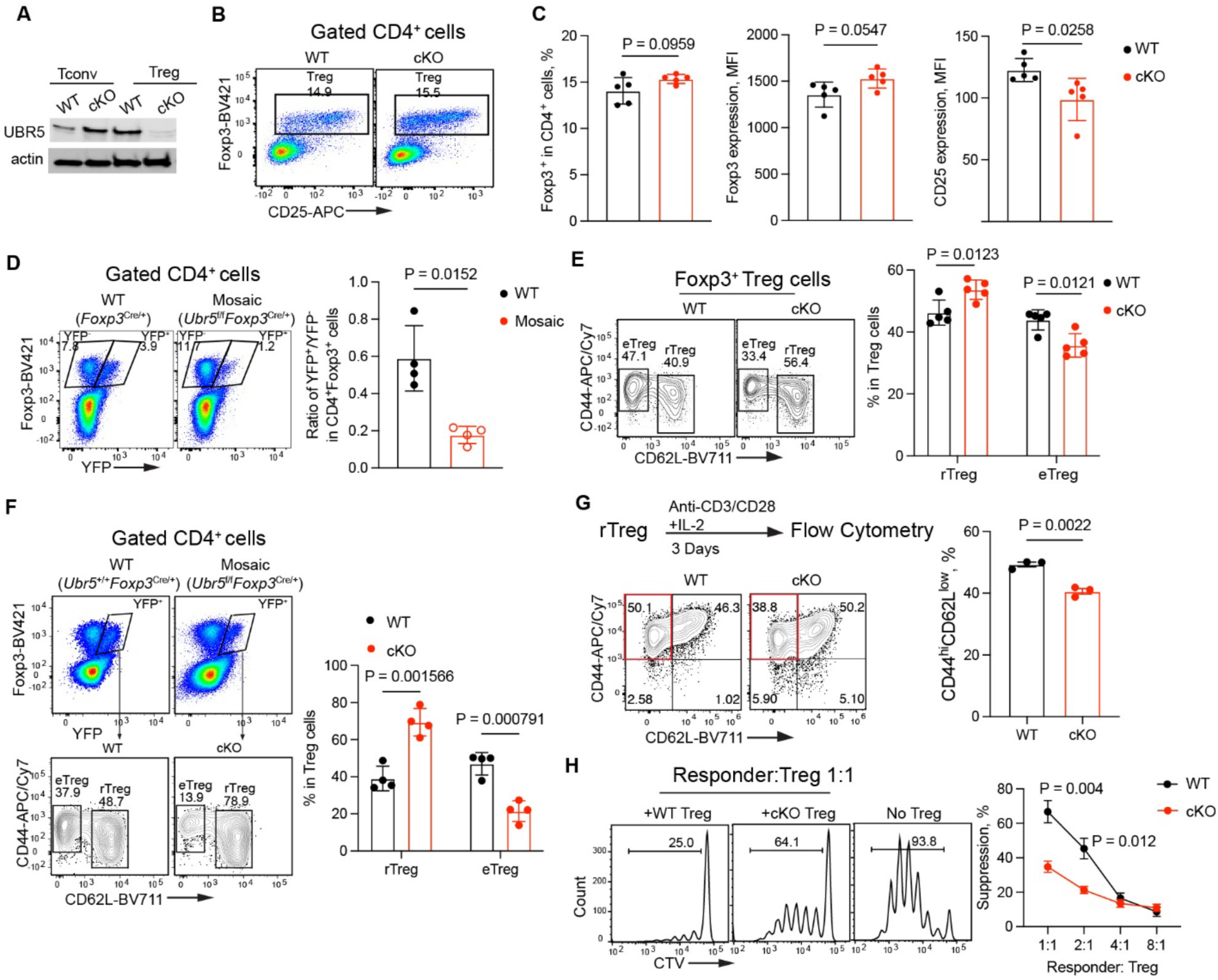
UBR5 intrinsically supports effector Treg differentiation and suppressive function. (A) Immunoblots of conventional T (Tconv) and Treg cells. (B) Representative flow cytometry plots show Treg cells from WT (*Foxp3*^Cre^) and UBR5 cKO (*Ubr5*^f/f^ *Foxp3*^Cre^) mice. Numbers indicate the percentage of gated cells. (C) Bar graphs show the percentage of Foxp3^+^ cells in CD4^+^ T cell populations (left), Foxp3 expression level (middle), and CD25 expression (right). MFI, median fluorescence intensity. (D) Representative plots of YFP^+^ and YFP^−^ Treg cells in CD4^+^ T cells from spleens of WT (*Foxp3*^Cre/+^) and mosaic (*Ubr5*^f/f^ *Foxp3*^Cre/+^) mice (left), and the ratio of YFP^+^/YFP^−^ Treg cells (right). Numbers indicate percentage in gating. (E) flow cytometry plots of CD44 and CD62L expression in Treg cells from WT and cKO mice (left), and the percentages of rTregs and eTregs (right). (F) CD4^+^ T cells from spleens of control or mosaic mice were analyzed by flow cytometry. YFP^+^ cells were further gated to analyze the expression of CD44 and CD62L (left). The percentage of eTregs in the YFP^+^ cell population is shown in the bar graph (right). (G) FACS-sorted rTreg cells were stimulated with anti-CD3/CD28 Abs and IL-2 for 3 days, then were analyzed for the expression of CD44 and CD62L. (H) CTV-labelled responder T cells were cultured with anti-CD3/CD28 Dynabeads and Tregs from WT and UBR5 cKO mice for 3 days (N = 3 mice).Cell proliferation was analyzed by flow cytometry. The representative histogram of responder proliferation (left) and quantification of suppressive activity of Treg cells (right). N = 4-5 mice/group from (A) to (F). Data are mean ± SD. Student’s two-tailed t test was used. Results are representative of two independent experiments.

Despite the loss of UBR5, Foxp3⁺ cell frequency and Foxp3 protein expression did not differ significantly (Figure 1C). Although CD25 expression was slightly reduced, IL-2R signaling—assessed by pSTAT5 induction—was intact (Figure S2A), indicating that UBR5 is dispensable for maintaining Treg identity under steady-state conditions.

To test the intrinsic role of UBR5 in Tregs, we generated *Ubr5*^f/f^ *Foxp3*^Cre/+^ “mosaic” female mice, which possessed both UBR5 KO Tregs (YFP^+^) and WT Tregs (YFP^−^) due to random inactivation of the X chromosome carrying the Foxp3-Cre allele in females. The ratio of UBR5-deficient and WT Treg populations in the mosaic mice was strongly reduced compared with that in *Foxp3*^Cre/+^ WT control mice (Figure 1D), indicating that UBR5 supports the competitive fitness of Tregs.

We next examined resting Treg (rTreg) and effector Treg (eTreg) subsets, which are distinguished by CD62L and CD44 expression. UBR5 cKO mice showed a significant reduction in eTregs with a corresponding increase in rTregs (Figure 1E). Mosaic mice confirmed a cell-intrinsic requirement for UBR5 in eTreg formation: UBR5-deficient YFP⁺ Tregs displayed an approximately 50% lower eTreg frequency than WT Tregs (Figure 1F). In vitro, UBR5-deficient rTregs showed impaired differentiation into eTregs in response to TCR stimulation (Figure 1G).

UBR5-deficient Tregs were significantly less effective at inhibiting naïve T-cell proliferation (Figure 1H). To exclude the possibility that reduced suppression reflected altered eTreg/rTreg subset composition, we assessed suppressive activity in sorted eTregs. UBR5-deficient eTregs exhibited reduced suppressive activity (Figure S2B), suggesting that UBR5 loss compromised the suppressive activity of eTregs. Consistent with compromised suppressive function, UBR5 cKO mice displayed increased CD44^hi^CD62L^lo^ effector-memory CD4⁺ T cells and a trend toward increased effector-memory CD8⁺ T cells in the spleen (Figure S2D–E). Thus, UBR5 is intrinsically required for eTreg differentiation and suppressive function.

### UBR5 maintains Treg proliferative fitness

We asked why total Treg frequency remained unchanged despite fewer eTregs in UBR5 cKO mice. It has been shown that eTregs rapidly die whereas rTregs proliferate and replenish eTreg pools (Miyara et al., 2009). Thus, peripheral Treg frequency is the net effect of concurrent proliferation and cell death. Because UBR5 deficiency promotes apoptosis in embryonic tissues and tumor cells (Saunders et al., 2004; Yu et al., 2024), we examined Treg viability. eTregs were more apoptotic than rTregs in the spleen of WT mice as expected (Figure 2A). Surprisingly, UBR5 cKO Tregs displayed reduced early apoptosis—most dramatically in eTregs (4-fold reduction; Figure 2B). Thus, apoptosis did not account for the reduced eTreg frequency at this time point.

**Figure 2.**
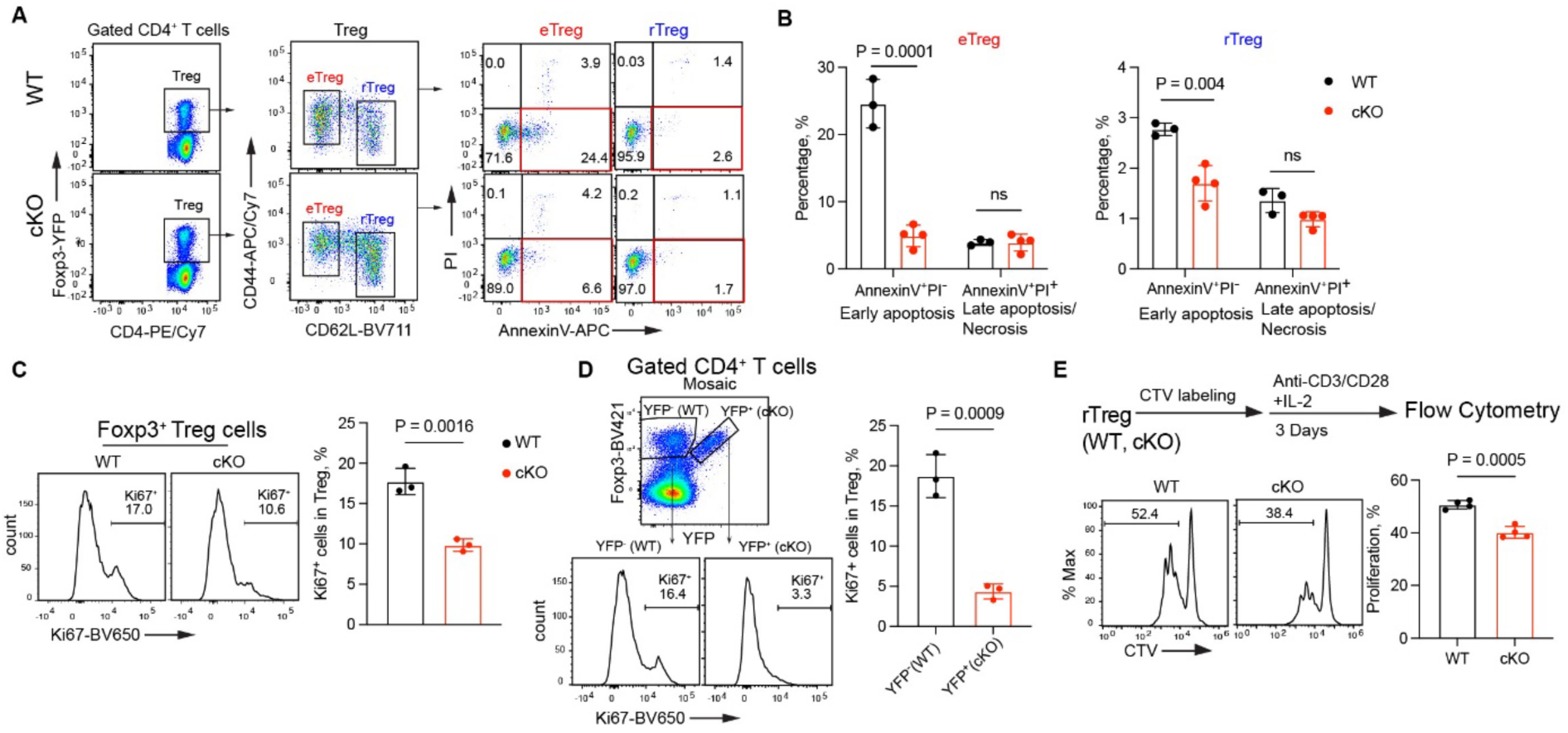
UBR5 maintains Treg proliferative fitness. (A) Splenocytes from WT and UBR5 cKO mice at 6 weeks of age were analyzed by flow cytometry. Treg cells were first gated as CD4^+^YFP^+^ cells (left), then further separated as eTregs and rTregs based on the expression of CD62L and CD44 (middle). eTregs and rTregs in apoptosis or necrosis were distinguished by the Annexin V and PI staining (right). Numbers indicate the percentage of cells in gating. (B) Percentages of cells in early apoptosis (Annexin V^+^PI^−^) and in late apoptosis/necrosis (Annexin V^+^PI^+^). (C) Ki67 expression in WT and cKO Tregs. (D) Splenocytes from mosaic (*Ubr5*^f/f^*Foxp3*^Cre/+^) female mice at 6 weeks of age were stained with antibodies against CD4, Foxp3 and Ki67. Treg cells were gated as Foxp3^+^, separated as YFP^−^ (WT) Treg and YFP^+^ (KO) Tregs (Top), and assessed for Ki67 expression (bottom). The percentages of Ki67^+^ cells in WT and cKO Tregs are shown (right). Two-tailed paired Student’s t test was used. (E) FACS-sorted rTreg cells were labeled with CTV, stimulated with anti-CD3/CD28 plus IL-2 for 3 days and analyzed by flow cytometry. Representative flow plots show cell proliferation indicated by CTV dilution. Bar graph shows the percentage of cells in proliferation. Data are mean ± SD. Student’s two-tailed t test was used in (B, C and E). Results are representative of two independent experiments.

We next examined Treg proliferation. Ki67⁺ Tregs were markedly reduced in UBR5 cKO mice (Figure 2C), a defect that was amplified in mosaic mice, where KO Tregs contained only ∼25% as many Ki67⁺ cells as WT Tregs (Figure 2D). To further investigate proliferation, we labeled rTreg cells with CellTrace Violet (CTV) dye and stimulated them with anti-CD3/CD28 plus IL-2. UBR5-deficient Tregs exhibited reduced proliferation (Figure 2E) compared with WT Tregs, indicating that UBR5 is required for maintaining Treg proliferation.

In summary, UBR5 loss impairs Treg proliferation, but concurrently reduces cell death, which may account for the absence of a significant change in total Treg frequency.

### UBR5 deficiency impairs effector-associated transcriptional programs and activates p53 signaling

To identify transcriptional mechanisms underlying the impaired eTreg compartment, we performed RNA-seq on WT, heterozygous, and UBR5-deficient rTregs and eTregs. Principal component analysis (PCA) confirmed clear segregation by both genotype and activation status (Figure 3A). UBR5 loss modestly altered rTregs (88 up- and 370 down-regulated genes) but profoundly altered eTregs (355 up- and 1,006 down-regulated genes) (Figure 3B). Gene set enrichment analysis (GSEA) revealed that genes associated with oxidative phosphorylation were enriched, while genes associated with “MTORC1 signaling”, “IL-2-STAT5 signaling”, “Interferon gamma response”, “inflammatory response” and “interferon alpha response” were underrepresented in UBR5 cKO Tregs (Figure 3C), consistent with the effector differentiation defects observed in UBR5-deficient Tregs. Indeed, genes for effector and suppressive function (e.g. *Gzmb, Nt5e, Entpd1, Fgl2, Lag3, Pdcd1, and Tigit*) and genes for migration and adhesion (e.g. *Cxcr6, Cxcr4, Cxcr5, Ccr6,* and *Itgae*) were down-regulated in UBR5-deficient Tregs (Figure 3D). GSEA also showed that effector-up genes (Dias et al., 2017) were underrepresented in UBR5-deficient Tregs (Figure 3E).

**Figure 3.**
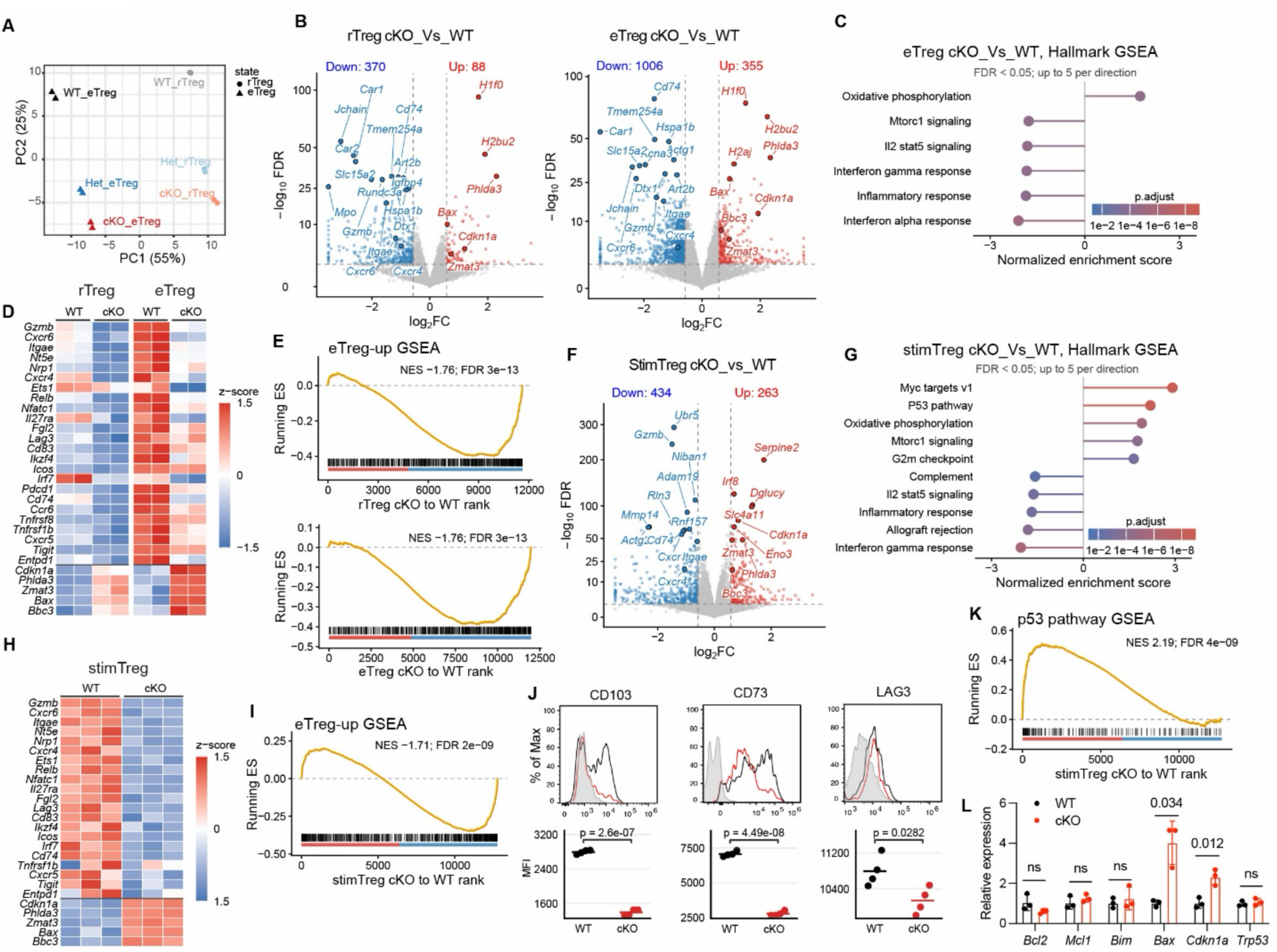
UBR5 deficiency impairs effector-associated transcriptional programs and activates p53 signaling. (A) Principal component analysis (PCA) of the RNA-seq data. CD44^hi^CD62^low^ eTregs and CD44^low^CD62^hi^ rTregs were sorted from UBR5 WT (*Foxp3*^Cre^), UBR5 heterozygous (Het, *Ubr5*^f/+^*Foxp3*^Cre^) and UBR5 cKO (*Ubr5*^f/f^*Foxp3*^Cre^) mouse spleens (n = 2 mice per genotype) and subjected to RNA-seq. (B) Volcano plots of RNA-seq comparing the expression of genes in rTreg cKO versus WT and in eTreg cKO versus WT. Differentially expressed genes (DEGs) (fold change, 1.5-fold, adjusted p value [padj] < 0.01) are shown in red (Up) and blue (Down) and enumerated. Some representative genes are labeled. (C) Gene set enrichment analysis (GSEA) using Hallmark gene sets. Positive and negative normalized enrichment score indicate enrichment toward genes higher and lower in eTreg cKO, respectively. (D) Heatmap showing expression of selected effector-associated genes and five p53 targets in cKO versus WT Tregs. (E) GSEA comparing the relative expression of eTreg-up genes (Dias et al., 2017) in rTreg cKO versus WT (top) and eTreg cKO versus WT (bottom). NES, normalized enrichment score. (F) Volcano plot of RNA-seq comparing the expression of genes in stimulated (Stim) Treg cKO versus WT. Sorted Treg were activated with anti-CD3/CD28 plus IL-2 for 48 h and subjected to RNA-seq. (G) GSEA using Hallmark gene sets. Positive and negative normalized enrichment score indicate enrichment toward genes higher and lower in StimTreg cKO, respectively. (H) Heatmap showing expression of selected effector-associated genes and five p53 targets in Stim Treg cKO versus WT. (I) GSEA comparing the relative expression of eTreg-up genes (Dias et al., 2017) in Stim Treg cKO versus WT. (J) Flow cytometry analysis comparing the expression of effector genes in Stimu Tregs. Student’s two-tailed t test was used (N = 4 mice / genpotype). (K) GSEA comparing the relative expression of genes in p53 pathway in Stim Treg cKO versus WT. (L) qPCR analysis for apoptosis-related genes in Stim Tregs. Gene expression in cKO cells was normalized to expression in WT cells. Data are mean ± SEM. Student’s two-tailed t test was used (N = 3 mice/genotype).

Upon stimulation, 263 genes were upregulated and 434 genes were downregulated in stimulated UBR5 cKO Tregs compared with WT Tregs (Figure 3F). Genes related to “MYC targets”, “p53 pathway” and oxidative phosphorylation were enriched, while genes related to “IL-2-STAT5 signaling”, “inflammatory response” and “Interferon gamma response” were underrepresented in stimulated UBR5 cKO Tregs (Figure 3G). Genes for effector and suppressive function were expressed at lower levels in UBR5 cKO Tregs compared with WT Tregs (Figure 3H, 3I). Flow cytometry analysis confirmed that UBR5 cKO Tregs expressed lower levels of CD103 (encoded by *Itgae*), CD73 (encoded by *Nt5e*) and LAG3 than WT Tregs (Figure 3J).

Notably, canonical p53 target genes (*Phlda3*, *Zmat3*, *Cdkn1a/p21*) (Fischer, 2017) were strongly upregulated in rTregs, eTregs, and stimulated Tregs (Figure 3B, 3D and 3H). GSEA revealed that genes in the p53 pathway were strongly enriched in UBR5 cKO Tregs (Figure 3K). qPCR analysis showed that loss of UBR5 significantly increased the expression of canonical p53 target genes *Bax* and *Cdkn1a* without affecting the expression of *Trp53*, anti-apoptotic genes *Bcl-2* and *Mcl-1,* and pro-apoptotic gene *Bim* (Figure 3L).

In summary, UBR5 loss selectively dismantles the effector Treg program while upregulating p53 target genes.

### Treg-specific UBR5 deletion enhances antitumor immunity and responsiveness to PD-1 blockade

Since eTregs are the predominant Treg subset in tumors (Sakaguchi et al., 2020), we tested whether UBR5 loss in Tregs alters antitumor immunity. In the MC38 colon adenocarcinoma model, UBR5 cKO mice exhibited markedly reduced tumor growth and significantly prolonged survival (Figure 4A–B). The number of tumor-infiltrating CD8^+^ T cells was significantly increased in UBR5 cKO mice, although the numbers of tumor-infiltrating CD4^+^ T cells and Tregs and the frequency of Foxp3^+^ Tregs were comparable between genotypes (Figure 4C). Effector function of CD4⁺ and CD8⁺ TILs were markedly enhanced in UBR5 cKO mice, as exemplified by enhanced production of IFN-γ in this condition (Figure 4D). Strikingly, Tregs in UBR5 cKO tumors also produced more IFN-γ (Figure 4D), a hallmark of Treg fragility associated with reduced suppressive activity (Overacre-Delgoffe et al., 2017). In the immunotherapy-refractory B16-F10 melanoma model, UBR5 cKO mice were sensitized to anti–PD-1 therapy, showing reduced tumor growth and improved survival (Figure 4E–F). These results show that UBR5 is required for maintaining functional, suppressive Tregs in the TME, and its loss unleashes robust antitumor immunity.

**Figure 4.**
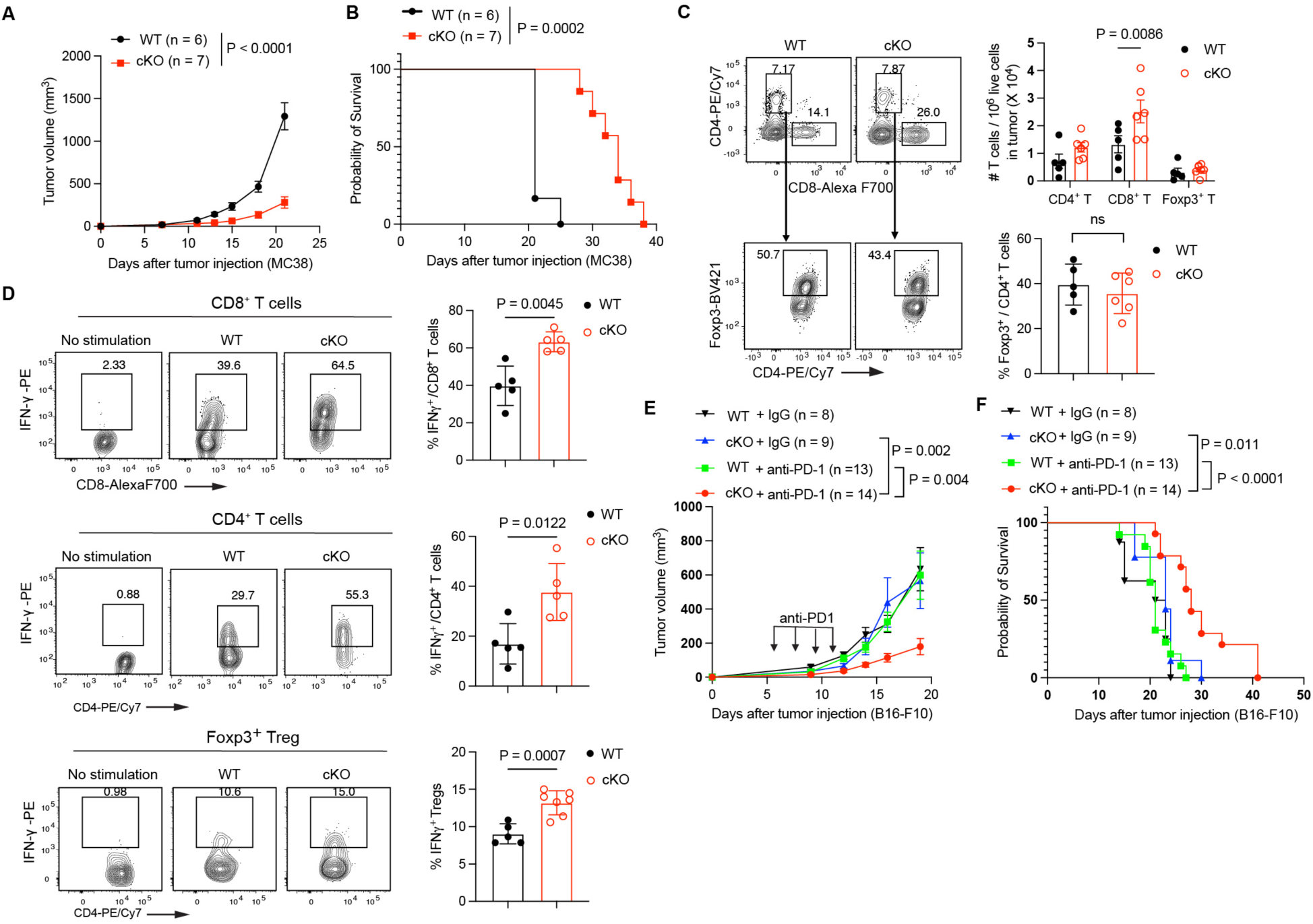
Treg-specific UBR5 deletion enhances antitumor immunity and responsiveness to PD-1 blockade. (A) MC38 tumor growth and (B) survival in WT and UBR5 cKO mice. (C) Tumor infiltrating lymphocytes (TILs) were analyzed on day 18 from MC38 tumors grown in WT and UBR5 cKO mice. Representative flow cytometry contour plots show CD4^+^ and CD8^+^ T cells in live CD45^+^ CD3^+^ tumor infiltrating cells (top). Cell numbers are shown in the bar graph. Foxp3^+^ Tregs were further gated from CD4^+^ T cells (bottom). The percentage of Foxp3+ cells is presented in bar graph. (D) TILs were isolated on day 18 from MC38 tumors grown in WT and UBR5 cKO mice and were activated with a stimulation cocktail with GolgiStop for 6 h. IFN-γ expression in CD8^+^-T, CD4^+^-T and Foxp3^+^ -Treg cells were analyzed by flow cytometry. Numbers indicate percentage in gating. (E) B16-F10 tumor growth and (F) survival in WT and UBR5 cKO mice. Anti-PD-1 or IgG isotype antibody was injected on days 6, 8, 10, and 12 after tumor inoculation as indicated by the arrows. N = 5 to 6 mice/group in (C, D). Data are mean ± SD in (C, D), or mean ± SEM (A, E). Two-way ANOVA followed by multiple comparisons test was used in (A, E). Log-rank test was used in (B) and (F). Mann–Whitney U tests were used in (C, D). Results are representative of two independent experiments.

### UBR5 sustains the intratumoral effector Treg program

To further define UBR5’s role in shaping tumor-infiltrating Tregs, we performed scRNA-seq analysis on CD4⁺ TILs from MC38 tumors. We identified naïve CD4⁺ T cells (Tn), Th1 cells, and distinct transcriptional states of Tregs—including intermediate, proliferative, and Gzmb⁺ Treg clusters (Figure 5A; Figure S3A). Treg identity was verified by *Foxp3*, *Il2ra*, and *Ctla4* expression (Figure S3B). Comparison of Tregs from WT and UBR5 cKO mice revealed that cKO Tregs had a modest reduction in the proportion of C2_Treg_Gzmb Tregs and an increase in C5_Treg_Intermediate (Figure 5B, Figure S3C).

**Figure 5.**
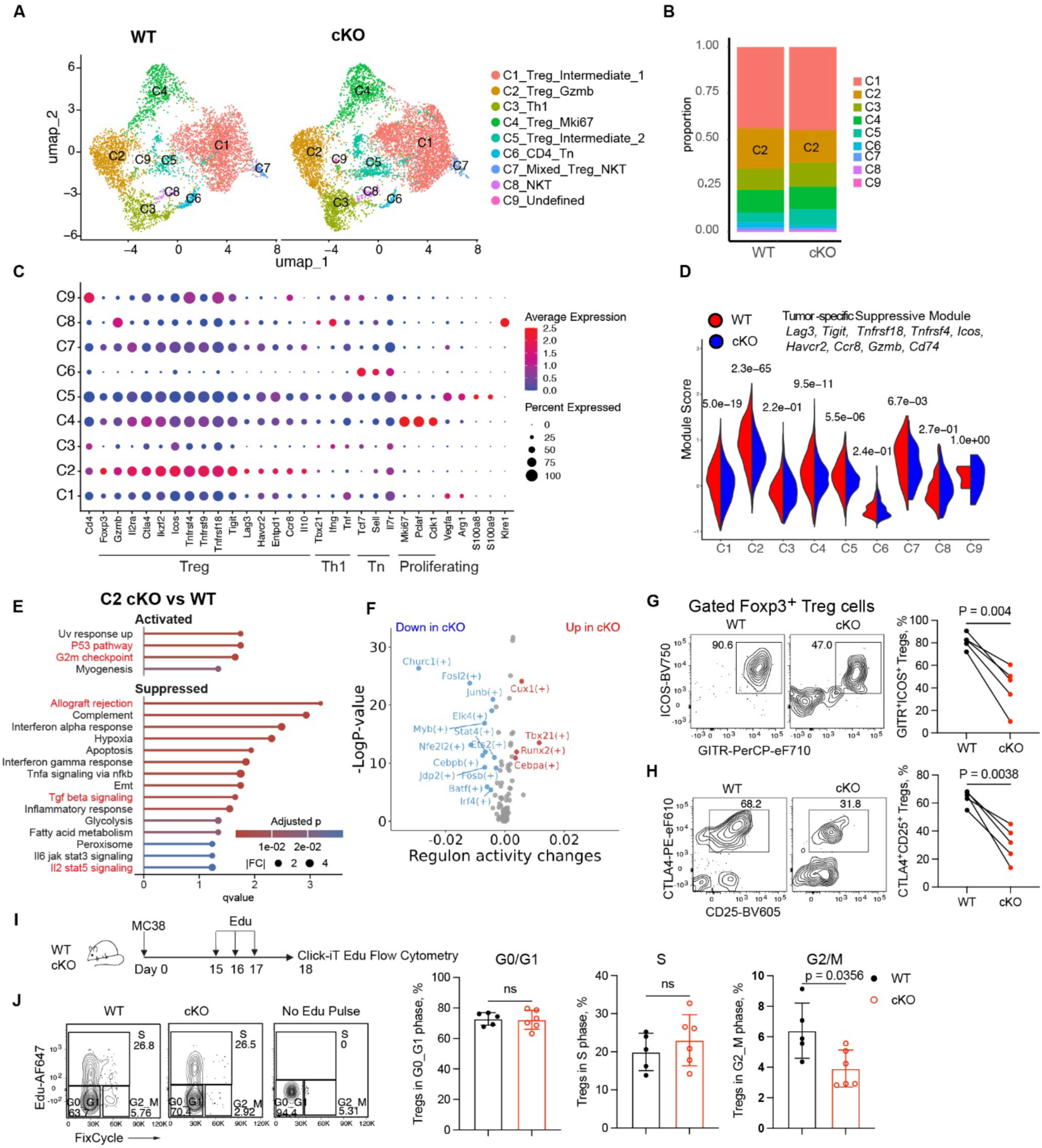
UBR5 sustains the intratumoral effector Treg program. (A) UMAP of CD4^+^ T cells. Tumor infiltrating lymphocytes (TILs) were enriched by 36% Percoll centrifugation and pooled from three mice per genotype. CD45^+^CD3^+^TCRb^+^CD4^+^ cells were FACS-sorted and used for scRNA-seq, yielding one library per genotype. Nine distinct clusters (C1-C9) were identified. (B) The proportion of each cluster. (C) Dot plot shows the canonical marker genes of each cluster. (D) Tumor specific module (TSM) scores comparing the 9 clusters from WT and cKO Tregs. (E) Activated and suppressed hallmark gene sets in C2_cKO versus WT Tregs (adjusted P value < 0.05). (F) Volcano plot showing the regulon activity changes in C2_cKO versus WT Tregs. The top altered regulons are highlighted. (G, H) MC38 tumor was inoculated into female mosaic mice. On day 18 after inoculation, tumor-infiltrating Treg cells were assessed by flow cytometry. WT Tregs from the CD4^+^ T compartment were gated as YFP^−^Foxp3^+^ and UBR5 cKO Tregs as YFP^+^Foxp3^+^. Effector Tregs were further gated as ICOS⁺GITR⁺ (G) or CTLA4⁺CD25⁺ subsets (H). The percentages of each subset are shown in the dot plots on the right. Two-tailed paired Student’s t test was used (N = 5 mice/group). (I) Experimental design to examine cell cycling in tumor infiltrating Tregs. (J) Flow cytometry plots of cell cycling in tumor Tregs from WT or UBR5 cKO mice and the percentage of Tregs in G0/G1 phase, S phase and G2/M phase. Data are mean ± SD. Student’s two-tailed t test was used (N = 5-6 mice/group). Results are representative of two independent experiments.

Tregs in Cluster 1 (C1) highly expressed *Arg1* and *Vegfa* (Figure 5C), a characteristic of Tregs under hypoxic conditions (Facciabene et al., 2011; Kempkes et al., 2019). C2 cells were tumor eTregs, exhibiting relatively high expression of Ikzf2*, Ccr8, Il2ra, Icos, Tnfrsf4, Tnfrsf9, Tnfrsf18, Ctla4, Lag3, Tigit* and *Gzmb* (Figure 5C). Tregs in C4 were actively proliferating, reflected by high expression of the cell-cycle markers *Mki67, Pclaf, Cdk1* (Figure 5C). C5 cells expressed high levels of *S100a8* and *S100a9* (Figure 5C), which are expressed in nTreg cells (Palmer et al., 2019; Stubbington et al., 2015). Tregs in C4 and C5 from UBR5 cKO mice expressed more *Ifng* than the counterparts from WT mice (Figure S3D), consistent with a shift toward an inflammatory phenotype.

Analysis of DEGs in intratumoral eTregs (C2_Treg_Gzmb) showed that UBR5 KO Tregs displayed a less effector phenotype, reflected by lower expression of *Gzmb, Cd74, Cxcr4, Lag3, Tigit, Klf2, Ccr8, Icos, among others* (Figure S3E). To further determine whether deletion of *UBR5* impairs tumor-specific Treg cell function, we created a tumor-associated suppressive module (TSM) based on the proposed tumor-specific eTreg signature genes (Bonnin et al., 2024; Chaudhuri et al., 2024), using AddModuleScore function in Seurat. We found that the TSM had the highest score in WT C2 (Figure 5D), which is consistent with its eTreg phenotype. While UBR5-deficient Tregs displayed lower tumor-associated suppressive module scores across Treg subsets, with the largest difference observed in the C2 effector Treg cluster (Figure 5D). Pathway analysis (Bibby et al., 2022) showed strong activation of p53 and G2/M checkpoint pathways in cKO Tregs and suppression of allograft rejection, interferon-α response, hypoxia adaptation, and cytokine-signaling pathways (Figure 5E).

A SCENIC regulon analysis (Aibar et al., 2017) of the C2 tumor-infiltrating Treg cluster revealed that UBR5 deficiency was associated with a coordinated reduction in inferred activity in several transcriptional regulators that are well-established drivers of effector Treg differentiation (Figure 5F). Regulons governed by BATF and IRF4, which together form the core TCR-dependent activation module required for effector Treg programming in tumors (Alvisi et al., 2020; Itahashi et al., 2022; Vasanthakumar et al., 2015), showed significant reductions in UBR5 cKO C2 cells. The MYB regulon, a key determinant of effector Treg proliferation and maturation (Dias et al., 2017), was also markedly diminished, consistent with the impaired cycling and reduced effector phenotype of UBR5-deficient Tregs. In addition, the JunB regulon, which supports Treg functional stability and activation in inflammatory environments (Koizumi et al., 2018), showed decreased activity in cKO cells. In contrast, the Tbx21 regulon associated with Th1-adapted Treg programs (Dykema et al., 2023) showed increased activity in KO Tregs. Consistent with this shift, UBR5 cKO Tregs in clusters 4 and 5 displayed increased *Ifng* expression (Figure S3D). To validate the defects of tumor-infiltrating UBR5-deficient Tregs, we used flow cytometry to analyze the tumor eTreg subsets and cell cycling. UBR5-deficient Tregs displayed decreased ICOS⁺GITR⁺ and CTLA4⁺CD25⁺ subsets (Figure 5G, 5H) compared with WT Tregs, consistent with impaired acquisition of effector markers in vivo in the TME.

Because “p53 pathway” and “G2M checkpoint” were activated in UBR5-deficient Tregs, we analyzed cell cycling and proliferation in tumor-infiltrating Tregs by pulsing tumor-bearing mice with 5-ethynyl-2’-deoxyuridine (EdU) (Figure 5I). UBR5-deficient Tregs showed a significant loss of G2/M-phase cells compared with WT counterparts (Figure 5J), consistent with altered cell-cycle progression.

Together, these data indicate that UBR5 is required to maintain the transcription factor network that supports tumor eTreg differentiation.

### Activation of the p53–p21 checkpoint mediates key defects in UBR5-deficient Tregs

Within tumors, UBR5-deficient Tregs exhibited an altered eTreg program (Figure 3E, 3I, Figure 5D) and showed altered cell-cycle distribution, with a selective loss of cells in G2/M (Figure 5J). Pathway analysis implicated p53 as a potential cause. UBR5 loss consistently elevated several canonical p53 target genes without upregulating Trp53 mRNA (Figure 3L), consistent with post-transcriptional regulation of p53 protein. Furthermore, downregulation of p53 is required for antigen-specific CD4^+^ T-cell proliferation (Watanabe et al., 2014). We therefore asked whether p53 is responsible for the defects in UBR5-deficient Tregs.

We generated Treg-conditional p53 knockout (p53 cKO) and UBR5/p53 double knockout (cDKO) mice. Interestingly, p53 cKO Tregs expressed more UBR5 protein, suggesting a possible feedback relationship between UBR5 and p53 (Figure 6A). UBR5/p53 cDKO abolished p21 expression (Figure 6A), consistent with regulation of the p53-p21 pathway. p53 deletion restored eTreg frequencies to WT levels in conditional knockout mice (Figure 6B) and in mosaic mice (Figure S4A). p53 deletion increased Treg proliferation and rescued the proliferative defect of UBR5-deficient Tregs (Figure 6C). p53 deletion restored the expression of eTreg markers (e.g. CD103, CD73) in UBR5-deficient Tregs (Figure 6D). Functionally, p53 deletion restored tumor growth in UBR5 cKO mice, as cDKO mice displayed MC38 tumor growth comparable to WT mice (Figure 6E). p53 deletion abolished the enhanced CD8^+^ TIL infiltration in UBR5 cKO mice (Figure 6F). EdU-labeling revealed that deletion of p53 rescued cell-cycle progression of UBR5-deficient Tregs in TME, reflected by increased G2/M population in cDKO mice (Figure 6G-H). Thus, UBR5 supports Treg cell cycling, proliferation and effector phenotype by restraining p53 activation.

**Figure 6.**
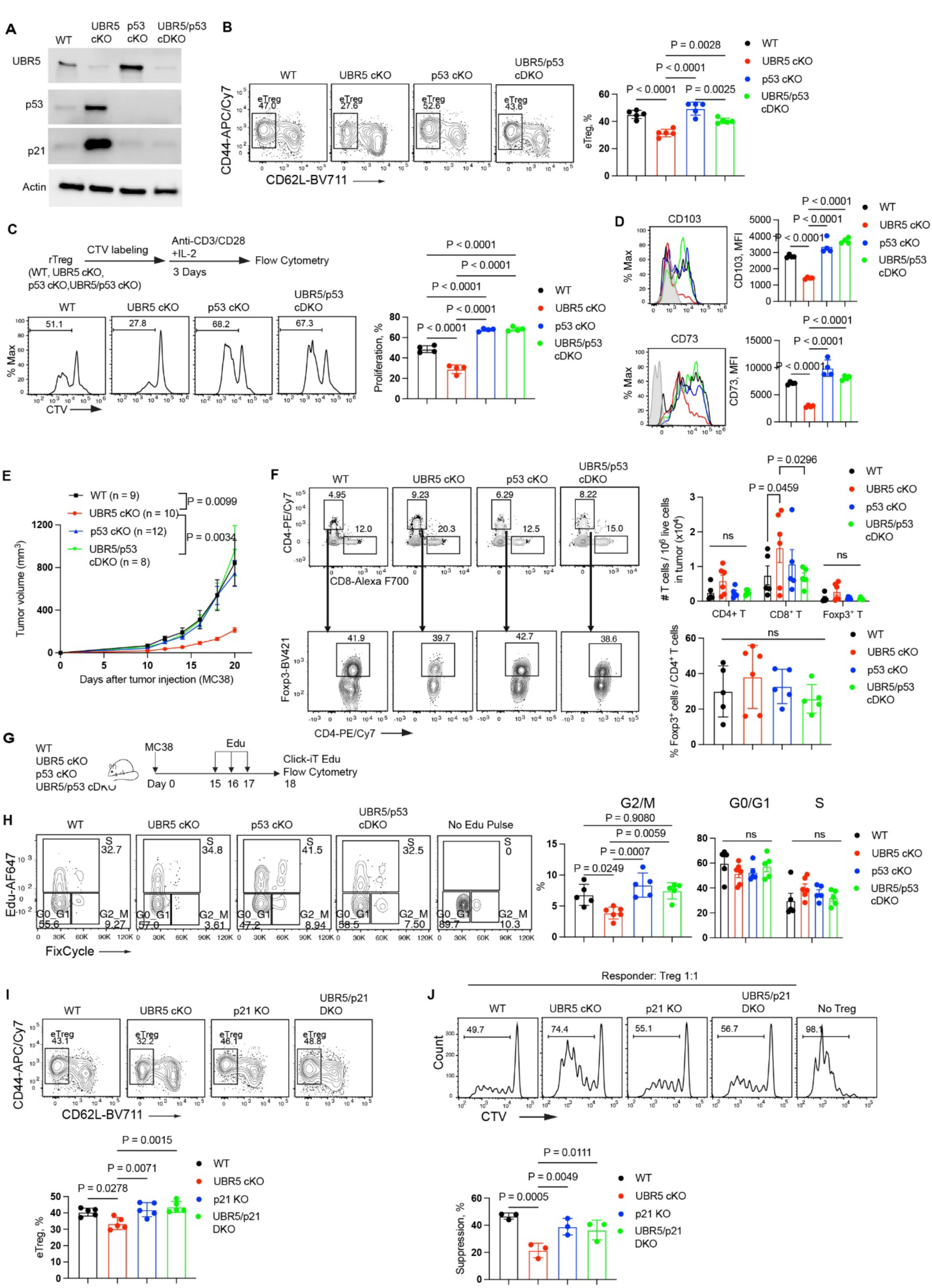
Activation of the p53–p21 checkpoint mediates key defects in UBR5-deficient Tregs. (A) Immunoblots of WT-, UBR5 cKO-, p53 cKO-, and p53/UBR5 cDKO-Tregs stimulated with anti-CD3/CD28 and IL-2 for 48 h. (B) Treg cells from WT, UBR5 cKO, p53 cKO and UBR5/p53 cDKO mice were analyzed for CD44 and CD62L expression (left) and the percentages of aTregs are shown in bar graph (right). (C) Representative flow plots show cell proliferation indicated by CTV dilution. Bar graph shows the percentage of cells in proliferation. (D) Flow cytometry analysis comparing the CD103 and CD73 expression in WT-, UBR5 cKO-, p53 cKO-, and p53/UBR5 cDKO-Tregs from (C). (E) MC38 tumor growth in WT, UBR5 cKO, p53 cKO and UBR5/p53 cDKO mice. Data are pooled from two experiments and shown as mean ± SEM. Two-way ANOVA followed by multiple comparisons test was used (N = 8-12 mice/group). (F) Tumor infiltrating lymphocytes (TILs) were analyzed on day 18 from MC38 tumors grown in WT, UBR5 cKO-, p53 cKO-, and p53/UBR5 cDKO mice. Representative flow cytometry contour plots show CD4^+^ and CD8^+^ T cells in live CD45^+^ CD3^+^ tumor infiltrating cells (top). Cell numbers are shown in the bar graph. Foxp3^+^ Tregs were further gated from CD4^+^ T cells (bottom). The percentage of Foxp3^+^ cells is presented in bar graph. (G) Experimental design to examine cell cycling in tumor infiltrating Tregs. (H) Flow cytometry plots of cell cycling in tumor Tregs from WT, UBR5 cKO, p53 cKO, and UBR5/p53 cDKO mice and the percentages of Tregs in G0_G1 phase, S phase and G2_M phase. N = 5-6 mice/group. (I) Treg cells from WT, UBR5 cKO, p21 KO and UBR5/p21 DKO mice were analyzed for CD44 and CD62L expression (top) and the percentages of eTregs are shown in bar graph (bottom). N = 5 mice/group. (J) CTV-labelled Responder cells were co-cultured with Treg cells from WT, UBR5 cKO, p21 KO, or UBR5/p21 DKO mice at a ratio of 1:1 and stimulated with anti-CD3/anti-CD28 beads for 3 days. Cell proliferation was analyzed by flow cytometry. The representative histogram of responder proliferation (top). Quantification of suppressive activity of Treg cells is shown in bar graph (bottom). One-way ANOVA followed by multiple comparisons was used in (B-D, F-J). Results are representative of two independent experiments.

We next tested whether p21 mediates this effect downstream of p53. Deletion of p21 restored eTreg frequencies in conditional knockout mice (Figure 6I) and in mosaic mice (Figure S4B) to WT levels. Because global deletion of p21 may induce T cell exhaustion in tumors (Thoma et al., 2024), our p21/UBR5 DKO mice may not be appropriate for testing Treg function in vivo in tumor models. We thus isolated p21/UBR5 DKO Tregs and performed an in vitro suppression assay. Deletion of p21 rescued the suppressive activity of UBR5 cKO Treg cells (Figure 6J). Thus, elevated p21 downstream of stabilized p53 constrains Treg suppressive function in UBR5-deficient Tregs.

In summary, activation of p53-p21 checkpoint after UBR5 loss compromises Treg cell cycling, proliferation and function.

### UBR5 promotes p53 protein turnover through its HECT domain

Next we asked whether UBR5 is involved in p53 ubiquitination and proteasomal degradation, a major mechanism controlling p53 protein stability (Kruse and Gu, 2009). Unstimulated WT Tregs lacked detectable p53 and its downstream target p21, whereas UBR5 cKO Tregs accumulated basal p53 and showed marked accumulation of p53 and p21 upon stimulation with anti-CD3/CD28 and IL-2 (Figure 7A). Cycloheximide chase experiments with WT and UBR5 cKO Tregs showed that p53 half-life was prolonged in UBR5 cKO Tregs (∼80 min vs < 20 min for WT p53 in Tregs) (Figure 7B). Treatment of Tregs with the proteasome inhibitor MG-132 increased p53 protein levels in both WT and UBR5-deficient cells (Figure 7C), indicating that UBR5 may not be the only E3 ligase mediating the proteasome-dependent degradation of p53 in Tregs. Similarly, the abundance of MDM2 protein, which forms a negative feedback loop with p53, also increased after MG-132 treatment. Loss of UBR5 reduced endogenous p53 poly-ubiquitination by half (Figure 7D). These results associate UBR5 loss with reduced p53 ubiquitination.

**Figure 7.**
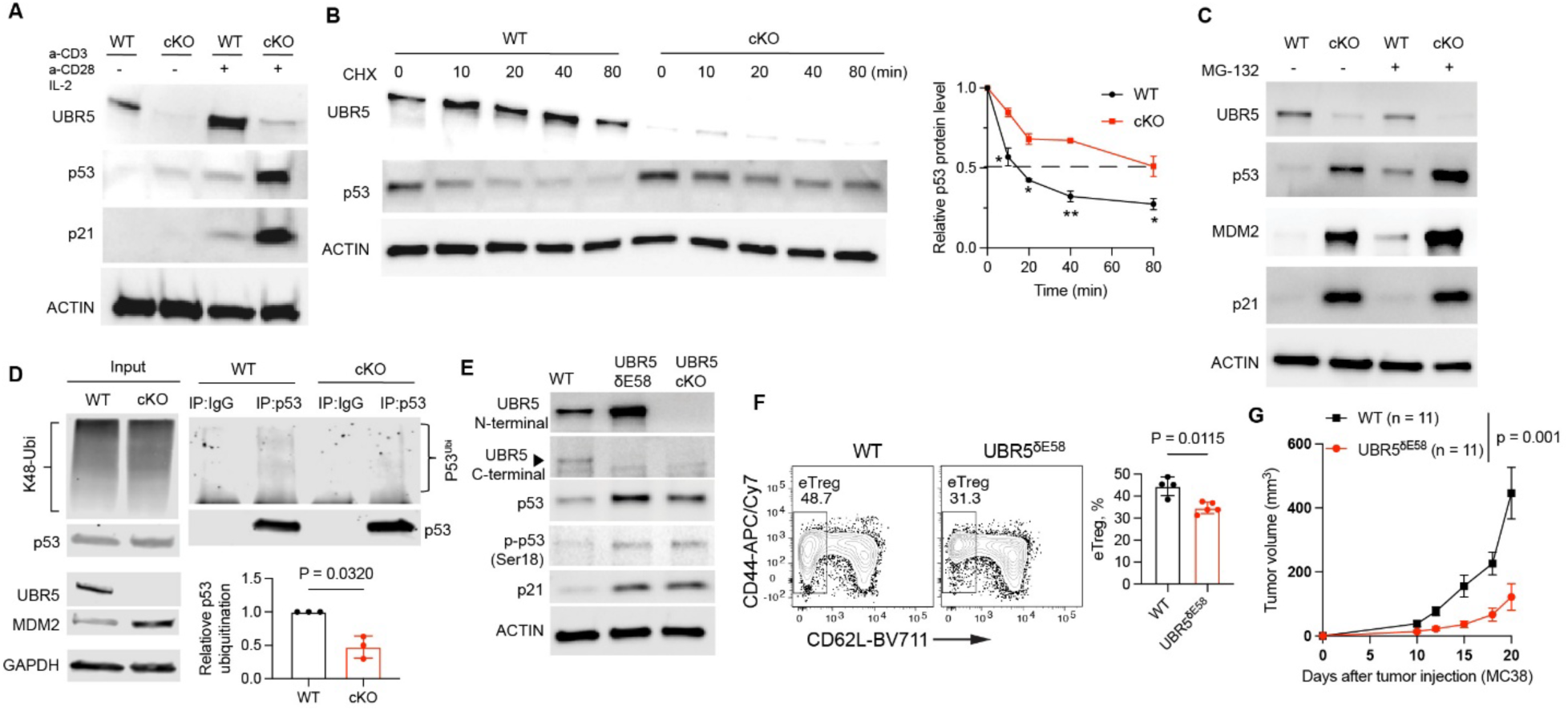
UBR5 promotes p53 protein turnover through its HECT domain. (A) Immunoblots of WT and UBR5 cKO Tregs stimulated with (+) or without (−) anti-CD3/CD28 and IL-2 for 48 h. (B) WT and UBR5 cKO Tregs were stimulated with anti-CD3/28 for 48 h, then treated with cycloheximide (CHX) for indicated times. Immunoblotting was performed to quantify the decay rate of p53 protein. P53 protein abundance was normalized to actin in each sample. Relative p53 expression at each time point was normalized to the expression at time zero for each genotype. Data are shown as mean± SEM of three independent experiments. Student’s two-tailed t test was used to compare relative p53 protein levels in WT and cKO Tregs at each time point. *, *p* < 0.05; **, *p* < 0.01. (C) WT and UBR5 cKO Tregs were treated without (−) or with (+) MG-132 (10 µM) and analyzed for UBR5, p53, MDM2 and p21 protein levels by immunoblotting. (D) Activated WT and UBR5 cKO cells were treated with MG132 for 2 h and analyzed for K48-linked polyubiquitination (K48-Ubi), UBR5, p53 and MDM2 protein levels by immunoblotting (left). Endogenous p53 was immunoprecipitated (IP) from cell lysates using anti-p53 antibody. K48-Ubi of p53 (p53^Ubi^) and total p53 was examined by immunoblotting with anti-K48 ubiquitin antibody (top-right panel). p53^Ubi^ was normalized to immunoprecipitated total p53 in each sample. Relative p53 ubiquitination in cKO cells was normalized to p53 ubiquitination in WT cells. (E) Immunoblots of WT-, UBR5^δE58^- and UBR5 cKO-Tregs stimulated with anti-CD3/CD28 and IL-2 for 48 h. (F) Flow cytometry plots of CD44 and CD62L expression in Treg cells from WT and UBR5^δE58^ mice (left), and the percentage of eTregs (right). Data are mean ± SD. Student’s two-tailed t test was used (N = 4-5 mice/group). (G) MC38 tumor growth in WT and UBR5^δE58^ mice; Data are mean ± SEM. Two-way ANOVA followed by multiple comparisons test was used (N = 11 mice/group). Results are representative of two independent experiments.

Moreover, we asked whether the E3 ubiquitin ligase activity of UBR5 contributes to regulation of p53 in Tregs. UBR5’s E3 ligase activity is mediated through its Homologous to the E6-AP Carboxyl Terminus (HECT) domain in the C-terminus (Wang et al., 2023). We crossed *Foxp3*^Cre^ mice with the UBR5 HECT domain floxed (*HECT*^f/f^) mice (Swenson et al., 2020), which allows conditional deletion of exon 58 in the HECT domain leading to a truncated UBR5 protein that lacks the cysteine at residue 2768 required for ubiquitin conjugation (referred to as UBR5^δE58^). In UBR5^δE58^ Tregs, elevated p53, p-p53^Ser18^ and p21 were observed, which phenocopied the full UBR5 cKO Tregs (Figure 7E). In UBR5^δE58^ mice, eTreg frequency was significantly reduced relative to control mice (Figure 7F), and MC38 tumors grew more slowly than in controls (Figure 7G), recapitulating the tumor control conferred by Treg-specific UBR5 deletion.

Together, these results support a role for the UBR5 E3 ubiquitin ligase activity in controlling the p53 protein level in Tregs.

### UBR5 expression is associated with human tumor eTreg features and immunotherapy nonresponse

We queried UBR5 using the Cancer Immune Data Engine (Gong et al., 2025), which integrates bulk-tumor transcriptomic and clinical-outcome datasets. Bulk-tumor UBR5 expression showed generally positive risk-association scores across the Immune Therapy and TCGA cohorts, consistent with an association between higher UBR5 expression and adverse clinical outcome (Figure 8A). However, this analysis is not cell-type specific; consequently, the observed associations may reflect UBR5 expression in cancer cells, Tregs, other immune populations, or a combination of these compartments.

**Figure 8.**
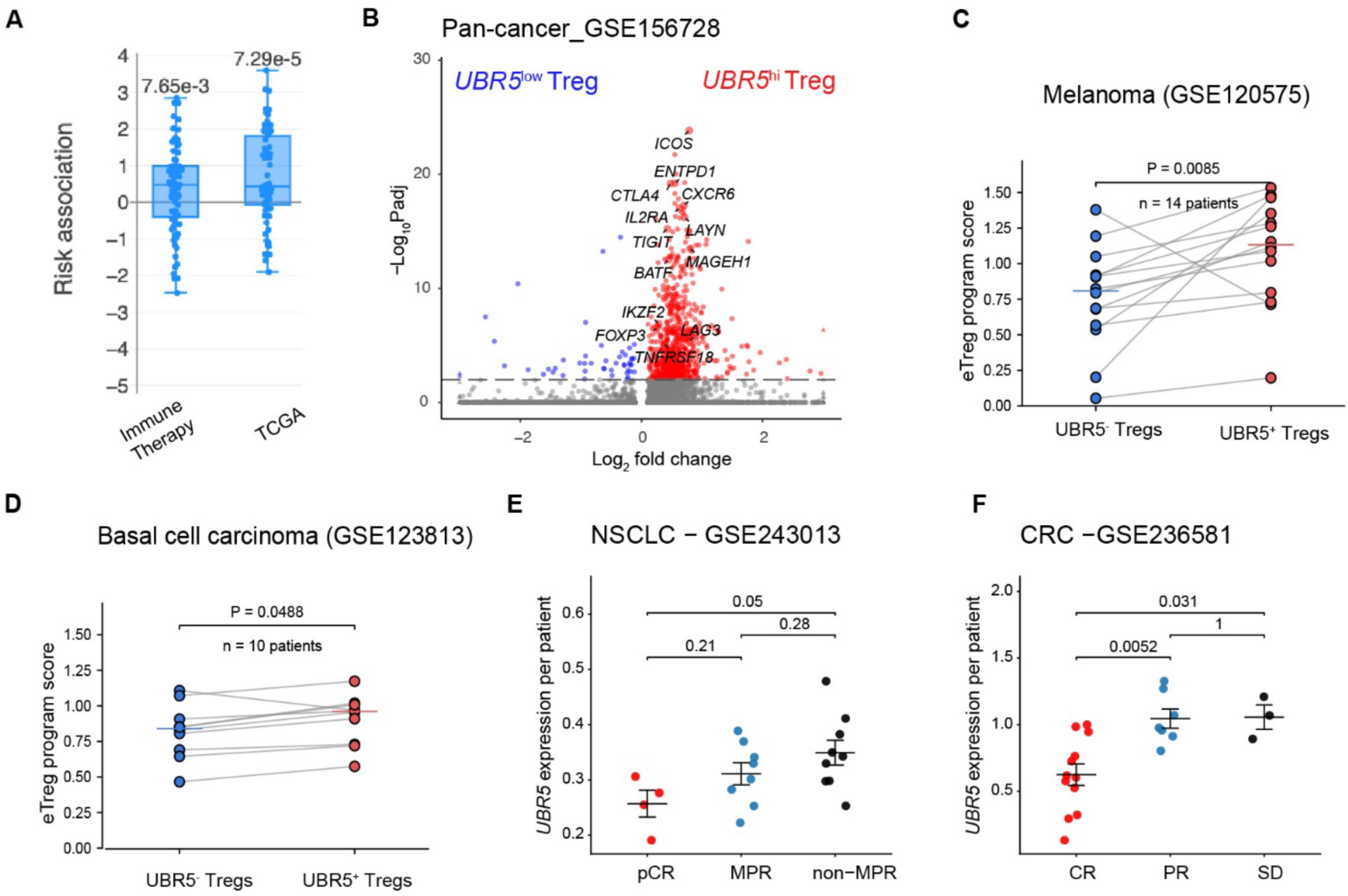
UBR5 marks a conserved human eTreg state and is associated with immune checkpoint blockade (ICB) nonresponse. (A) Risk association between *UBR5* mRNA expression and clinical outcomes in Immune Therapy and TCGA cohorts. Plot was generated using Cancer Immune Data Engine. Each dot represents one clinical cohort. p values were computed using the two-sided Wilcoxon signed-rank test comparing group values and zero. (B) Volcano plot of differentially expressed genes within *UBR5*^hi^ versus *UBR5*^low^ human tumor infiltrating Tregs in pan-cancer single-cell T-cell atlas. (C-D) Patient-level comparison of sequencing-depth-normalized eTreg program scores between UBR5⁺ and UBR5⁻ Tregs in pretreatment melanoma and basal cell carcinoma cohorts, respectively. Each line connects the mean scores of the two Treg populations within the same patient. Horizontal bars indicate cohort medians. P values were determined using two-sided paired Wilcoxon signed-rank tests. (E–F) Patient-level mean *UBR5* expression in tumor-infiltrating Tregs from post-treatment NSCLC (E) and colorectal cancer (F) samples, stratified by clinical response to anti–PD-1 therapy. Statistical comparisons were performed using pairwise Wilcoxon tests. pCR, pathological complete response (pCR); MPR, major pathological response MPR; CR, complete response; PR, partial response; SD, stable disease.

To further assess the relevance of UBR5 to human tumor immunity, we examined its expression in tumor-infiltrating Tregs from a pan-cancer single-cell T-cell atlas encompassing 21 malignancies (Zheng et al., 2021). Compared with UBR5^low^ Tregs, UBR5^hi^ Tregs displayed an effector-like transcriptional program characterized by increased expression of *FOXP3, IKZF2, IL2RA, ICOS, CTLA4, TIGIT, BATF,* and *MYB*, markers associated with activated tumor eTregs (Figure 8B).

To determine whether UBR5 identifies an activated human tumor-Treg state independently of clinical-response classification, we performed patient-aware analyses of two public immune checkpoint blockade (ICB) pretreatment single-cell cohorts. Within each patient, UBR5 transcript-detectable Tregs exhibited higher eTreg program scores than UBR5-undetectable Tregs in melanoma (Sade-Feldman et al., 2018) (median paired difference = 0.276; P = 0.0085) and basal cell carcinoma (Yost et al., 2019) (median paired difference = 0.103; P = 0.0488) (Figure 8C-D). Thus, the association between UBR5 expression and an eTreg transcriptional state was reproduced across two human tumor types.

We next compared UBR5 expression in Tregs from responders versus nonresponders to anti–PD-1 monotherapy in non-small cell lung cancer (NSCLC) (Liu et al., 2025) and colorectal cancer (CRC) (Chen et al., 2024). Complete responders (CR) exhibited lower UBR5 expression than non-responders (Figure 8E–F).

Together, these human data associate UBR5 with an activated tumor-eTreg state and with ICB nonresponse in selected cohorts.

## Discussion

Regulatory T cell (Treg) accumulation within tumors represents a major barrier to effective antitumor immunity (Shang et al., 2015), yet the mechanisms that sustain the Treg effector program and suppressive stability under chronic activation in the TME remain incompletely defined. Here, we identify the E3 ubiquitin ligase UBR5 as an important regulator that enables tumor-infiltrating Tregs to maintain proliferative capacity, effector differentiation, and suppressive function. Using conditional and mosaic genetic models, we show that UBR5 acts intrinsically in Tregs to support Treg effector programs. Loss of UBR5 leads to impaired Treg suppression and enhanced antitumor immunity. Our work provides two major lines of evidence that UBR5 is required for the Treg effector program. First, UBR5-deficient Tregs display cell-intrinsic defects in eTreg differentiation, cycling, and suppressive function. These defects are apparent within the TME, where Tregs rely on rapid proliferation to maintain effector programs. Second, transcriptomic and single-cell analyses showed that UBR5 loss alters the intratumoral Treg transcriptional landscape—downregulating hallmark eTreg genes while upregulating p53 target genes linked to cell-cycle arrest. These findings establish UBR5 as a key molecular node that couples cell-cycle control to eTreg differentiation in cancer.

Functionally, Treg-specific deletion of UBR5 unleashes antitumor immunity in two distinct tumor models. In the MC38 model, UBR5 loss in Tregs alone was sufficient to suppress tumor growth. In the immunotherapy-resistant B16-F10 melanoma model, UBR5 loss in Tregs sensitized tumors to anti–PD-1 therapy. Increased IFN-γ production by CD4⁺ and CD8⁺ tumor-infiltrating lymphocytes and the emergence of IFN-γ⁺ ‘fragile’ Tregs reflect a shift from a suppressive to a proinflammatory milieu (Kang and Zappasodi, 2023). These findings highlight UBR5 as a resistance node that protects Treg suppressive fitness and limits effector responses under PD-1 pathway inhibition.

An important insight from this study is the identification of the p53–p21 axis as a pathway restrained by UBR5 in Tregs. UBR5-deficient Tregs accumulated p53 and p21 proteins despite unchanged p53 mRNA levels, implicating post-transcriptional regulation. Despite the accumulation of p53 and increased expression of its proapoptotic target Bax, UBR5-deficient eTregs exhibited reduced Annexin V positivity at steady state. This observation suggests that the consequences of p53 activation in these cells are manifested predominantly as impaired cell-cycle progression rather than increased detectable apoptosis. Reduced proliferative activity and altered activation states may also contribute to the lower frequency of apoptotic cells. Although our data do not exclude differences in apoptotic kinetics or clearance, the genetic rescue experiments identify p53–p21-mediated proliferative restraint as a major contributor to the impaired fitness of UBR5-deficient Tregs. These findings reveal that UBR5 antagonizes the p53–p21 checkpoint to enable cell cycling needed for Treg suppressive function, especially within tumors where Tregs are exposed to hypoxia, oxidative and metabolic stress that trigger cell-cycle stress.

How UBR5 restrains p53 protein abundance depends on the cellular context. Smits found that UBR5 knockdown results in elevated p53 protein abundance both in transformed and untransformed cells without significantly affecting p53 mRNA abundance or p53 protein half-life, and speculated that UBR5 regulates p53 by participating in the microRNA-mediated translational silencing (Smits, 2012). However, the E3 ubiquitin ligase activity of UBR5 is not required for miRNA silencing (Su et al., 2011). Saez et al. reported that loss of UBR5 led to abnormal accumulation of ribosomal RNA processing intermediates, which triggered “nucleolar stress” in embryonic stem cells (Saez et al., 2020). During nucleolar stress, assembly factors and ribosomal proteins bind to MDM2. This binding blocks MDM2-p53 interaction, which results in the stabilization and accumulation of p53 protein. Ling et al. showed that silencing UBR5 induced p53-Ser15 phosphorylation, a modification that stabilizes p53 by reducing its binding to MDM2 (Ling and Lin, 2011). UBR5 binds p53 and sterically blocks Ser15 phosphorylation by ataxia telangiectasia mutated (ATM). However, this blocking does not require the HECT domain of UBR5. We observed that loss of full-length UBR5 (UBR5 cKO) and HECT domain (UBR5^δE58^) equally elevated p53-Ser18 (equivalent to human p53-Ser15) and total p53 protein, suggesting that the HECT domain is required for UBR5 to control p53 in Tregs.

The function of p53 in Tregs has been controversial. Deletion of p53 using CD4-Cre reduced Tregs in autoimmune and inflammatory disease models (Kawashima et al., 2013; Zhang et al., 2023a). In contrast, activation of p53 using an inhibitor of the MDM2/MDM4 heterodimer decreased Tregs in peripheral blood mononuclear cells from patients with thyroid cancer (Arena et al., 2021). Recent studies demonstrate that p53 supports Treg function through ALDH6A1-dependent propionyl-CoA metabolism and histone propionylation (Xie et al., 2026), and preserves Foxp3 expression during exposure to inflammatory Th17 cytokines (Silveria et al., 2026). These findings establish beneficial functions of p53 that must be considered alongside its capacity to restrain cell expansion. Our genetic experiments show that p53 contributes to the defects caused by UBR5 deficiency. Furthermore, p53 deletion did not change the expression of Foxp3 in tumor-infiltrating Tregs (Figure 6D). The apparently divergent functions of p53 in Tregs may reflect differences in the magnitude, duration, and cellular context of p53 activation. Basal or transient p53 activity may support metabolic homeostasis and lineage maintenance, whereas sustained p53 accumulation may impose proliferative constraints that compromise effector differentiation. Our findings suggest that UBR5 contributes to the regulation of this balance by limiting p53 protein accumulation and downstream p21 induction. Thus, rather than being uniformly beneficial or detrimental to Treg function, p53 may exert context-dependent effects that require precise regulation during Treg activation and adaptation to inflammatory environments.

Our observation that UBR5 deficiency reduces K48-linked polyubiquitination of endogenous p53, together with the prolonged p53 half-life and the requirement for the UBR5 HECT domain, supports a role for UBR5 in promoting proteasomal p53 turnover in Tregs. However, these findings do not distinguish direct ubiquitination of p53 by UBR5 from cooperation with MDM2 or indirect regulation of the cellular pathways controlling p53 stability. Defining the molecular relationship between UBR5 and MDM2 will be important for understanding how Tregs restrain p53 activation during sustained proliferative responses.

Regulon analysis suggests candidate transcriptional networks underlying the altered phenotype of UBR5-deficient eTregs. Reduced inferred BATF and IRF4 activity is consistent with changes in effector-associated target genes (Alvisi et al., 2020; Itahashi et al., 2022), while reduced MYB activity is consistent with impaired proliferative fitness (Dias et al., 2017) and reduced JunB activity with altered functional stability (Koizumi et al., 2018). Increased inferred Tbx21 activity is consistent with the observed inflammatory features. These associations provide a framework for interpreting the transcriptional changes.

Our analysis of human datasets reveals that UBR5 expression is enriched in tumor-infiltrating Tregs from nonresponders after anti–PD-1 therapy. These analyses remain correlative and do not establish UBR5 as a universal pretreatment biomarker of ICB response. Functional perturbation studies in primary human tumor Tregs will be required to determine whether UBR5 is causally necessary for human eTreg fitness and ICB resistance.

Beyond cell-intrinsic effects, we observed increased UBR5 levels in Tconv cells from UBR5 cKO mice (Figure 1A), suggesting compensatory activation or altered homeostasis in response to impaired Treg suppression. This raises the intriguing possibility that UBR5 coordinates opposing programs in Tregs and Tconv cells to maintain immune balance—an idea that warrants future investigation.

Together, our findings identify UBR5-dependent regulation of p53 protein homeostasis as a mechanism that sustains effector Treg proliferative fitness and suppressive function. Disruption of this pathway compromises the effector Treg program without overt loss of Foxp3 expression, revealing a means of uncoupling Treg lineage identity from functional competence. In tumors, this disruption enhances antitumor immunity and responsiveness to PD-1 blockade. Given the established tumor-intrinsic functions of UBR5 (Liao et al., 2017; Shearer et al., 2015; Song et al., 2020a; Song et al., 2020b), therapeutic targeting of this ubiquitin ligase may simultaneously compromise malignant-cell fitness and Treg-mediated immune suppression. Determining the extent to which tumor-associated Tregs can be preferentially targeted without compromising systemic immune tolerance will be an important next step.

## Materials and methods

### Ethics / Animal Approval Statement

All mouse experiments were approved by the Institutional Animal Care and Use Committee at Weill Cornell Medicine (protocol #0804-739A) and performed in accordance with institutional guidelines.

### Reagent and resource

Critical reagents including antibodies, chemicals, assay kits, oligonucleotides and software are listed in supplementary Table S1.

### Mice

*Ubr5* exon 6–7 floxed mice (*Ubr5*^f/f^; C57BL/6 background) were generated by Cyagen Biosciences. UBR5 HECT-domain floxed mice (*HECT*^f/f^) were obtained from Dr. Shannon Buckley (University of Utah). *Foxp3*^YFP-Cre^ (*Foxp3*^Cre^, stock #016959) and *Trp53*^f/f^ (stock #008462) mice were purchased from The Jackson Laboratory. *Cdkn1a*^−/−^ (p21 KO) mice were acquired as previously described (Liu et al., 2009). *Ubr5*^f/f^ mice were crossed with *Foxp3*^Cre^ mice to generate F1 mice, which were intercrossed to obtain *Ubr5*^f/f^ *Foxp3*^Cre^ (Treg-conditional UBR5 knockout; UBR5 cKO) and *Ubr5*^+/+^ *Foxp3*^Cre^ (WT) littermates. For some experiments, female *Ubr5*^f/f^ *Foxp3*^Cre/+^ mosaic mice were used. *Ubr5*^f/f^ *Foxp3*^Cre/Cre^ mice were further crossed with *Trp53*^f/f^ or *Cdkn1a*^−/−^ mice to generate UBR5/p53 double-knockout (UBR5/p53 cDKO) or UBR5/p21 DKO strains, respectively. Mice were used at 6–10 weeks of age, and littermates matched for age and sex were used whenever possible. All mice were housed in specific pathogen–free facilities at Weill Cornell Medicine, and all procedures were approved by the Institutional Animal Care and Use Committee.

### Tumor models

MC38 murine colon adenocarcinoma cells (a gift from Dr. Gregory Sonnenberg, Weill Cornell Medicine) were maintained in RPMI-1640 supplemented with 10% FBS. B16-F10 melanoma cells (ATCC) were grown in DMEM + 10% FBS. Tumor cells (1×10⁵ per mouse) were injected subcutaneously (MC38) or intradermally (B16-F10) into the right flank. Anti–PD-1 antibody was administered intraperitoneally on days 6, 8, 10, and 12. Tumors were measured using calipers, and volume was calculated as (length × width²)/2. Mice were euthanized if tumors exceeded 1.5 cm in diameter, ulcerated, or if animals showed signs of morbidity.

### Cell preparation and culture

Splenocytes were dispersed through 70-µm strainers and red blood cells were lysed using Hybri-Max RBC lysis buffer (Sigma). T cells were enriched using the Pan-T kit (Miltenyi). Cells were stained with viability dye and antibodies to CD3, CD4, CD8, CD25, CD44, and CD62L, and sorted on a BD Symphony S. Sorting strategies are shown in Supplementary Figure S5. Tregs were purified as CD4⁺CD25⁺YFP⁺ cells; naïve CD4⁺ T cells were sorted as CD4⁺CD25⁻CD44^low^CD62L^hi^. T cell culture medium consisted of RPMI-1640 with 10% FBS, penicillin–streptomycin (100 µg/mL each), 1× non-essential amino acids, and 50 µM β-mercaptoethanol.

### Tumor digestion

Tumors were minced and digested for 40 min at 37 °C in RPMI containing collagenase IV (1 mg/mL; Worthington Biochemical) and DNase I (0.1 mg/mL; Worthington Biochemical). Digests were filtered through 70-µm strainers, washed, and subjected to 36% Percoll centrifugation at 500 × g for 10 min. Red blood cells (RBCs) were removed by brief incubation in RBC lysis buffer. The single-cell suspension was used for downstream analysis or sorting.

### Flow cytometry

Dead cells were stained with Zombie UV Viability (BioLegend) and excluded for downstream analysis. Fc receptors were blocked using anti-CD16/CD32 for 10 min at room temperature. Surface marker staining was performed in PBS with 2% FBS and 0.1% sodium azide for 30 min at 4 °C. All the antibodies against surface markers were used at a 1:400 dilution. Intracellular Foxp3 staining used the Foxp3/Transcription Factor kit (eBioscience). For cytokine staining, cells were stimulated with Cell Stimulation Cocktail plus protein transport inhibitors (eBioscience) for 6 h before fixation. For phospho-STAT5 measurement, cells were starved for 1 h at 37 °C, stimulated with IL-2 for 20 min, fixed in 2% paraformaldehyde, permeabilized in ice-cold methanol, and stained with anti-pSTAT5. All the antibodies against intracellular proteins were used at 1:50 dilutions. Annexin-V apoptosis assays were conducted using APC-Annexin V and PI in Annexin V binding buffer (BD Biosciences). Data were acquired on a BD Symphony A5 and analyzed in FlowJo v10. Gating strategies are shown in Supplementary Figure S5.

### Cell cycle analysis

Mice were Intraperitoneally injected with EdU (20 mg/kg) every day for 3 days and were sacrificed 16 h after the last injection. Tumors were digested to prepare a single-cell suspension. Cells were stained with antibodies against surface markers, fixed and permeabilized using Fix/ perm buffer from Foxp3/Transcription Factor kit. Cells were then stained with anti-Foxp3-PE and resuspended in perm/wash buffer. EdU was detected using Click-iT EdU plus flow cytometry kit (Invitrogen) according to the protocol. Cells were resuspended in 500 ul perm/wash buffer containing FixCycle Violet dye and incubated for 30 min before flow cytometry analysis.

### Immunoprecipitation

Treg cells were sorted from spleens of WT and UBR5 cKO mice and stimulated with plate-bound anti-CD3 (1 µg/mL) and anti-CD28 (1 µg/mL) in the presence of IL-2 (500 U/mL) for 48 h. MG-132 (10 µM) was added for the final 2 h. Cells (5 × 10^6^ per genotype) were washed twice in ice-cold PBS and lysed for 30 min on ice in Pierce lysis buffer (Thermo Fisher Scientific) supplemented with protease and phosphatase inhibitor cocktails (Thermo Fisher Scientific). Lysates were cleared by centrifugation at 12,000 × g for 10 min at 4 °C, and protein concentration was determined by BCA assay (Thermo Fisher Scientific). An aliquot (10%) of each lysate was reserved as input. For immunoprecipitation, protein was pre-cleared with Protein A/G magnetic beads for 1 h at 4 °C and then incubated overnight at 4 °C with rotation with anti-p53 (Proteintech, 10442-1-AP), anti-UBR5 (Santa Cruz, sc-515494), anti-MDM2 (Cell Signaling, 51541), or an equivalent amount of matched isotype control IgG. Immune complexes were captured with Protein A/G beads for 2 h at 4 °C, washed three times with lysis buffer, and eluted by boiling in Laemmli sample buffer for 5 min. Inputs and immunoprecipitated proteins were resolved by SDS-PAGE and analyzed by immunoblotting. K48-linked polyubiquitination of p53 was examined by immunoblotting with an anti-K48 ubiquitin antibody (Cell Signaling Technology, 8081).

### Immunoblotting

Cells were lysed in RIPA buffer (Thermo Fisher Scientific) supplemented with protease inhibitors. Proteins were resolved on SDS–PAGE gels, transferred to PVDF membranes, and probed overnight at 4 °C with primary antibodies. After incubation with HRP-conjugated secondary antibodies, blots were developed using SuperSignal West Femto substrate (Thermo Fisher Scientific) and imaged on a Bio-Rad ChemiDoc system.

### Real-time RT-PCR

RNA was isolated using the RNeasy Micro Kit (Qiagen). cDNA was prepared using the High-Capacity cDNA Reverse Transcription Kit (Thermo Fisher Scientific). cDNA samples were analyzed using PowerUp™ SYBR™ Green Master Mix for qPCR (Thermo Fisher Scientific) with primers indicated in key resource table. Results were normalized to those of housekeeping gene Actin and are reported as 2^−(Delta^ ^Delta^ ^Ct)^.

### In vitro Treg suppression assay

Tregs were isolated from mouse spleens using regulatory T cell isolation kit (Miltenyi) and activated for 20 h with plate-bound anti-CD3 (1 µg/mL) and anti-CD28 (1 µg/mL) in T-cell medium containing IL-2 (500 U/mL). Naïve CD8⁺ responder T cells were isolated by magnetic separation (Miltenyi) and labeled with CellTrace Violet (Invitrogen). Responder cells (4×10⁴ per well) were cultured alone or with Tregs at indicated ratios in the presence of CD3/CD28 Dynabeads (Bead: cell ratio 2:1) and IL-2 (10 U/ml). After 3 days, responder cell proliferation was assessed by flow cytometry. Suppression (%) was calculated as: 100 × [1 – (proliferation with Tregs / proliferation alone)].

### RNA-seq and analysis

Tregs were sorted and RNA was isolated using the RNeasy Micro Kit (Qiagen). RNA quality was assessed on an Agilent BioAnalyzer. cDNA was prepared using the SMART-Seq v4 Ultra Low Input RNA kit (Takara) and libraries were constructed with Nextera XT (Illumina). Libraries were sequenced on an Illumina NovaSeq X Plus (paired-end 2×50 bp). Reads were aligned to mm39 using STAR with default parameters. Differential gene expression was performed using DESeq2, and pathway enrichment using clusterProfiler (Wu et al., 2021).

### scRNA-seq and analysis

CD45⁺TCRβ⁺CD4⁺ tumor-infiltrating cells were sorted and processed using the 10x Genomics Chromium GEM-X Single Cell 3’ kit. Libraries were sequenced on an Illumina NovaSeq X Plus. Single-cell sequencing FASTQ files were processed using Cell Ranger V3.1 (10x Genomics). Reads were aligned to the mouse genome mm39 from ENSEMBL GRCm39. Counts data were analyzed using R package Seurat 5.0. Cells expressing fewer than 200 or more than 6500 genes and cells that have more than 5% mitochondrial counts were removed. scGate was used to select the pure T cells (*Ptprc*+, Cd3e+, *Trac*+, *Adgre1*-, *Siglech*-, *Ms4a1*-) (Andreatta et al., 2022). Samples were then merged, and data were normalized with a scale factor of 10000. Principal component analysis (PCA) was performed on variable genes, and 30 principal components were used for the FindNeighbors function and Uniform Manifold Approximation and Projection (UMAP) construction. FindClusters was performed serially at different resolutions. A resolution of 0.5 was selected to avoid over-segmentation. Cell clusters were manually annotated based on known markers. Differential gene expression analysis comparing Tregs from KO mice and WT mice was performed using FindMarkers functions for each cell type. EnhancedVolcano R package was used for visualization. Module scores were calculated using AddModuleScore function. FetchData function was then used to extract module scores for each cell. A Wilcoxon test was used to compare module scores between different conditions within clusters. For gene set enrichment analysis, msigdbr package was used to obtain mouse gene sets, SCPA package was then used for pathway enrichment analysis (Bibby et al., 2022). Python implementation of the SCENIC algorithm (pyscenic) was used for gene regulatory analysis and calculating the activities of the regulons (Van de Sande et al., 2020).

### Analysis of Public scRNA-seq datasets

Risk association between *UBR5* expression and clinical outcomes was analyzed with Cancer Immunology Data Engine (CIDE) (Gong et al., 2025). Published scRNA-seq datasets were downloaded from Gene Expression Omnibus (GEO) and processed in Seurat, retaining the original cluster annotation where available. The pan-cancer T cell atlas (GSE156728) comprises 21 cancer types from 316 individuals (Zheng et al., 2021). Treg cells from this atlas were classified as *UBR5*^hi^ or *UBR5*^low^ by whether their UBR5 expression fell above or below the median across all Tregs. Differential expression (adjusted *P* < 0.05) between the two groups was assessed with FindMarkers function in Seurat. Model-based Analysis of Single-cell Transcriptomics (MAST) model was applied to adjust dataset variability. To test whether *UBR5* identifies an activated human tumor-Treg state independently of clinical-response classification, we performed patient-aware analyses of two public immune checkpoint blockade (ICB) pretreatment single-cell cohorts (Sade-Feldman et al., 2018; Yost et al., 2019). Tumor-infiltrating Tregs were stratified by *UBR5* transcript detection (*UBR5*^+^ vs *UBR5*^−^), and the mean effector Treg (eTreg) program score was computed separately for each group within each patient. The eTreg score was defined by *ICOS, TNFRSF4, TNFRSF9, BATF, IRF4, MYB, GZMB, LAG3, ENTPD1, CXCR6*, and *ITGAE*, excluding *UBR5* and any clinical-response labels. *UBR5* mRNA expression in Tregs was compared between responders and non-responders to anti-PD-1 therapy in two additional cohorts (Chen et al., 2024; Liu et al., 2025), using post-treatment samples only and patient-level mean expression.

### Statistical analysis

GraphPad Prism was used for statistical analyses unless otherwise indicated. Tumor growth was analyzed by two-way ANOVA with Bonferroni correction. Comparisons between two groups used unpaired or paired Student’s t-tests or Mann–Whitney U tests as appropriate. Multiplicity-adjusted one-way ANOVA was used for >2 group comparisons. Survival curves were compared by log-rank test. Data represent mean ± SD.

## Online supplemental material

Figs. S1–S5 show additional expression, flow cytometry, single-cell analysis, genetic-rescue, sorting, and gating data. Table S1 lists key resources.

## Data availability

Bulk RNA-seq and single-cell RNA-seq datasets generated in this study have been deposited in the Gene Expression Omnibus (GEO) under accession number GSE316259, GSE316260, GSE316261, and GSE316262. Publicly available datasets analyzed in this study are cited in the manuscript and listed in the Methods section. Processed data and analysis scripts used for single-cell transcriptomic and regulon analyses are available from the corresponding authors upon reasonable request.

## Acknowledgments

This work was supported by a DOD Breast Cancer Research Program Breakthrough Awards (W81XWH-21-1-0261) to X.M and HT9425261E177 to X.M and G.L; an NIH grant (1 R01 CA273716-01A1) to X.M and P.Z; an NIH grant (R01 CA276053) to G.L.

## Author contributions

A.X. designed and performed experiments, analyzed data, and drafted the manuscript.

H.Z. conducted biochemical and co-immunoprecipitation experiments.

E.X. performed human dataset analyses and immunotherapy response correlation studies.

T.Z. conducted single-cell RNA-seq processing and regulon analyses.

S.M.B. provided UBR5 HECT-domain mutant mice and conceptual input.

C.Y. and P.Z. contributed phospho-p53 analyses and p21 knockout models.

G.L. provided expertise on tumor models and experimental design.

R.Z. provided expertise in regulatory T cell biology and tumor immunology.

X.M. supervised the study, acquired funding, and co-wrote the manuscript.

All authors reviewed and approved the final manuscript.

## Abbreviations

cDKO: conditional double knockout
cKO: conditional knockout
CRC: colorectal cancer
CTV: CellTrace Violet
DEG: differentially expressed gene
EdU: 5-ethynyl-2′-deoxyuridine
eTreg: effector regulatory T cell
GEO: Gene Expression Omnibus
GSEA: gene set enrichment analysis
HECT: homologous to E6-AP carboxyl terminus
ICB: immune checkpoint blockade
IEL: intraepithelial lymphocyte
MFI: mean fluorescence intensity
NES: normalized enrichment score
NSCLC: non-small cell lung cancer
PCA: principal component analysis
rTreg: resting regulatory T cell
SCENIC: single-cell regulatory network inference and clustering
scRNA-seq: single-cell RNA sequencing
TCM: central memory T cell
Tconv: conventional T cell
TEM: effector memory T cell
TIL: tumor-infiltrating lymphocyte
TME: tumor microenvironment
Treg: regulatory T cell
UMAP: uniform manifold approximation and projection
YFP: yellow fluorescent protein

