## supplemental figures for "UBR5 sustains effector regulatory T cell fitness by restraining the p53 checkpoint"

Figure S1

A

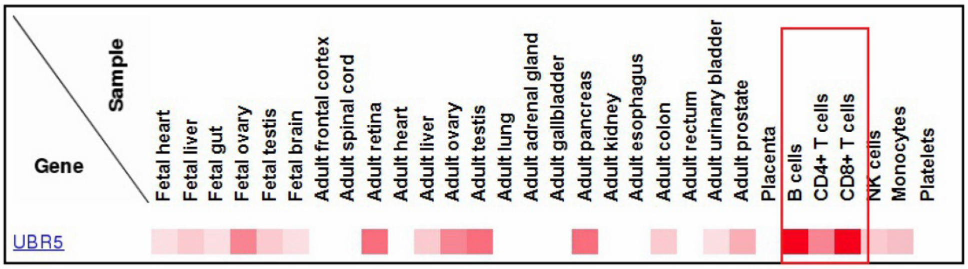
Sample
Gene
Fetal heart
Fetal liver
Fetal gut
Fetal ovary
Fetal testis
Fetal brain
Adult frontal cortex
Adult spinal cord
Adult retina
Adult heart
Adult liver
Adult ovary
Adult testis
Adult lung
Adult adrenal gland
Adult gallbladder
Adult pancreas
Adult kidney
Adult esophagus
Adult colon
Adult rectum
Adult urinary bladder
Adult prostate
Placenta
B cells
CD4+ T cells
CD8+ T cells
NK cells
Monocytes
Platelets
UBR5

B

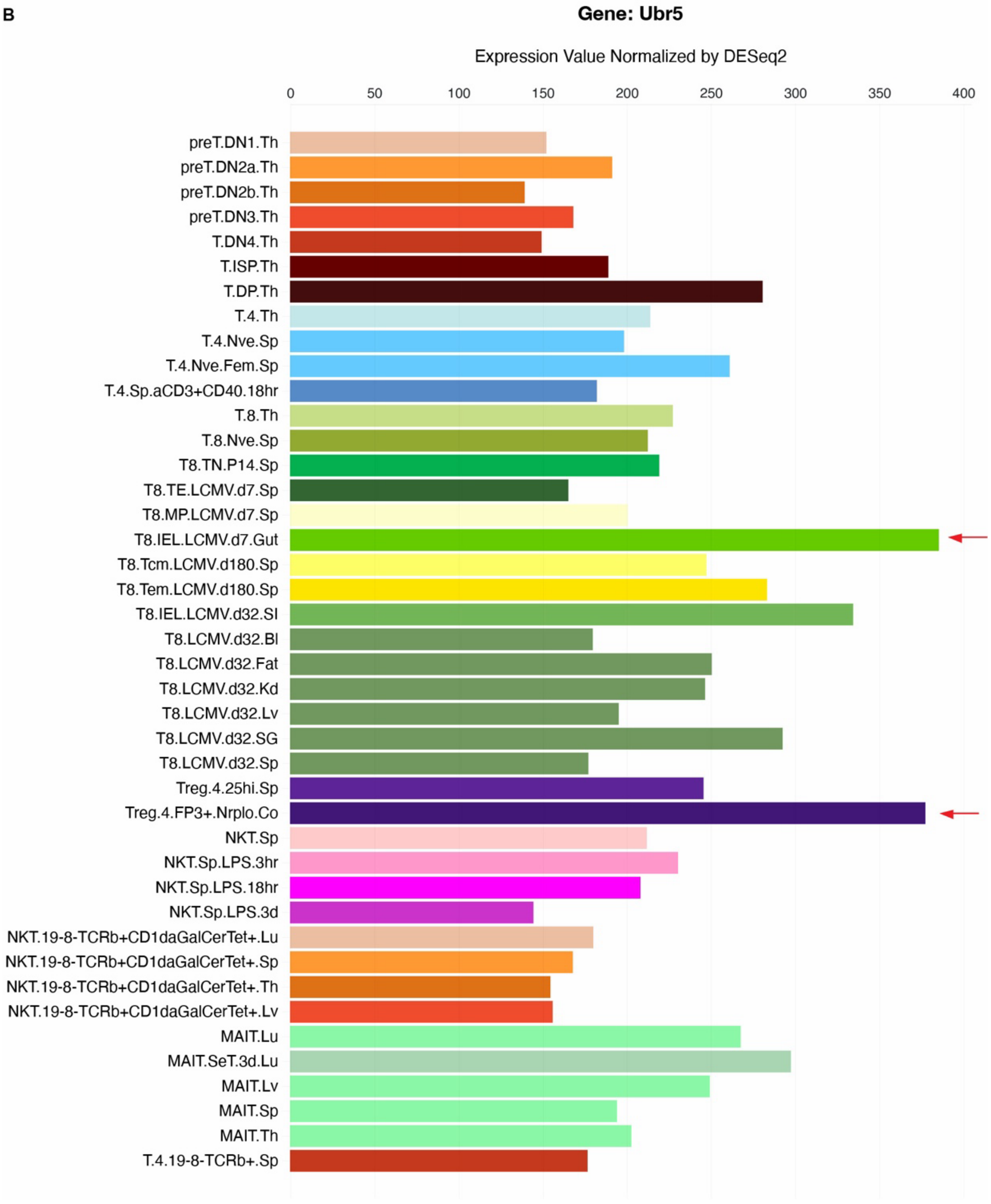
Gene: Ubr5
Expression Value Normalized by DESeq2
0
50
100
150
200
250
300
350
400
preT.DN1.Th
preT.DN2a.Th
preT.DN2b.Th
preT.DN3.Th
T.DN4.Th
T.ISP.Th
T.DP.Th
T.4.Th
T.4.Nve.Sp
T.4.Nve.Fem.Sp
T.4.Sp.aCD3+CD40.18hr
T.8.Th
T.8.Nve.Sp
T8.TN.P14.Sp
T8.TE.LCMV.d7.Sp
T8.MP.LCMV.d7.Sp
T8.IEL.LCMV.d7.Gut
T8.Tcm.LCMV.d180.Sp
T8.Tem.LCMV.d180.Sp
T8.IEL.LCMV.d32.SI
T8.LCMV.d32.Bl
T8.LCMV.d32.Fat
T8.LCMV.d32.Kd
T8.LCMV.d32.Lv
T8.LCMV.d32.SG
T8.LCMV.d32.Sp
Treg.4.25hi.Sp
Treg.4.FP3+.Nrplo.Co
NKT.Sp
NKT.Sp.LPS.3hr
NKT.Sp.LPS.18hr
NKT.Sp.LPS.3d
NKT.19-8-TCRb+CD1daGalCerTet+.Lu
NKT.19-8-TCRb+CD1daGalCerTet+.Sp
NKT.19-8-TCRb+CD1daGalCerTet+.Th
NKT.19-8-TCRb+CD1daGalCerTet+.Lv
MAIT.Lu
MAIT.SeT.3d.Lu
MAIT.Lv
MAIT.Sp
MAIT.Th
T.4.19-8-TCRb+.Sp

**Figure S1. UBR5 expression in T lymphocytes.** (A) UBR5 protein expression in human samples (http://humanproteomemap.org/). Red square highlights UBR5 expression in B and T lymphocytes. (B) UBR5 mRNA expression in mouse αβ T lymphocytes (https://www.immgen.org/). Red arrow points the expression of UBR5 in $CD8^+$ intraepithelial lymphocytes (IEL) in gut and $Foxp3^+Nrp^{lo}$ Treg cells in colon.

Figure S2

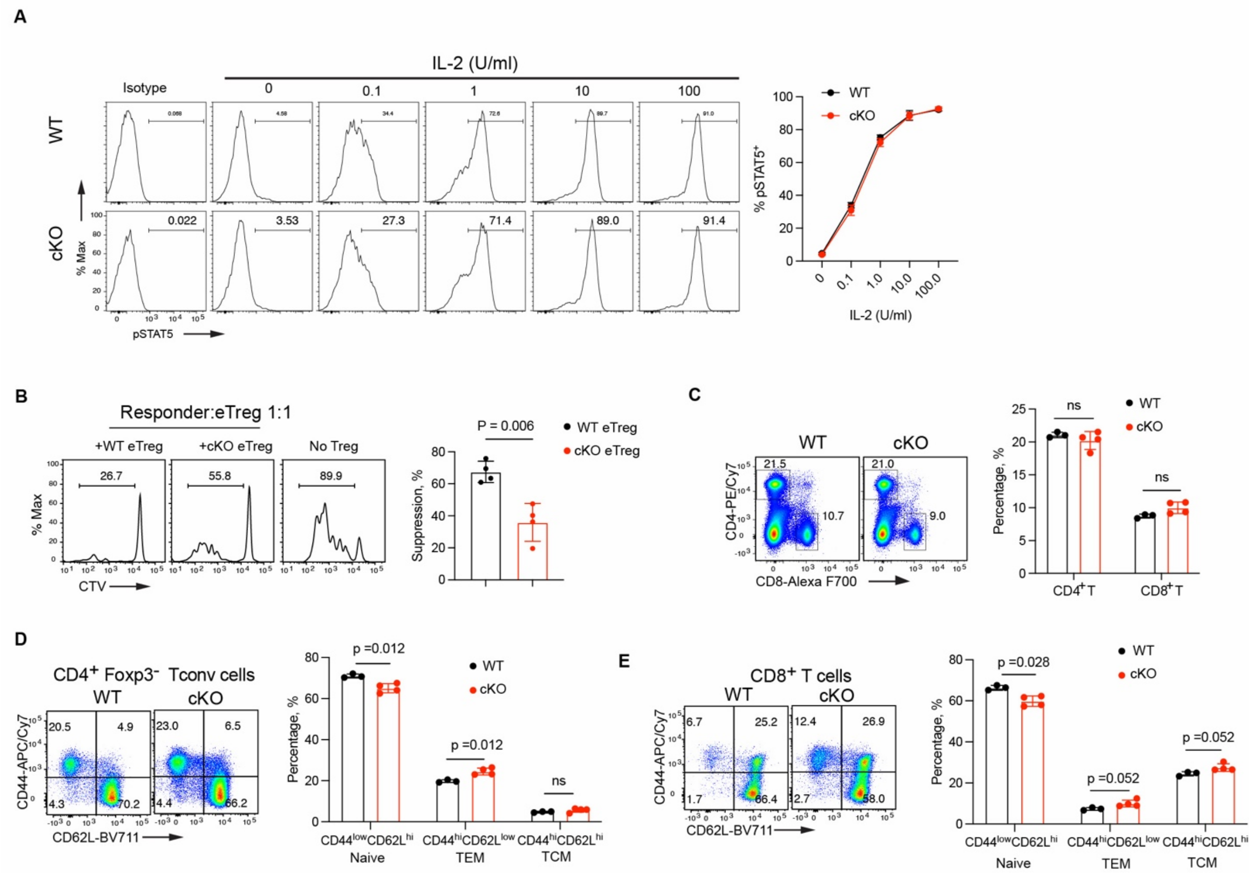


**Figure S2. UBR5 regulates eTreg function.** (A) Representative histogram plots (left) demonstrate induction of Stat5 phosphorylation upon IL-2 stimulation. Numbers indicate percentage (%) in gate. WT and UBR5 cKO Tregs were stimulated with grading doses of IL-2 for 20 min and then analyzed using flow cytometry. Line graph presents the percentage of pStat5 positive cells at different IL-2 concentration. (B) The representative histograms of responder cell proliferation (left) and quantification of suppressive activity of effectorTreg (eTreg) cells from WT and cKO mice (right). $CD44^{hi}CD62L^{low}$ eTreg cells were facs-sorted and co-cultured with CTV-labelled responder $CD8^{+}$ T cells and anti-CD3/CD28 Dynabeads for 3 days. Responder cell proliferation was then analyzed by flow cytometry. (C) Representative flow cytometry dot plots demonstrate $CD4^{+}$ and $CD8^{+}$ T cells from mice (left). Percentages of CD4+ and CD8+ T cells are shown in the bar graph (right). (D) Representative CD44 and CD62L expression in $CD4^{+}$ conventional T cells (Tconv) (left) from spleen, and percentages of naïve ($CD44^{low}CD62L^{hi}$), central memory T (TCM) ($CD44^{hi}CD62L^{hi}$), effector memory T (TEM) ($CD44^{hi}CD62^{low}$) in $CD4^{+}$ Tconv are quantified in bar graph (right). (E) Representative CD44 and CD62L expression in $CD8^{+}$

T cells (left) from spleen, and percentages of naïve, TCM ($CD44^{hi}CD62L^{hi}$), TEM cells ($CD44^{hi}CD62^{low}$) in $CD8^{+}$ T cells are quantified in bar graph (right). N = 4 mice/group from (C) to (E). Data are mean ± SD. Student's two-tailed t test was used. Results are representative of two independent experiments.

Figure S3

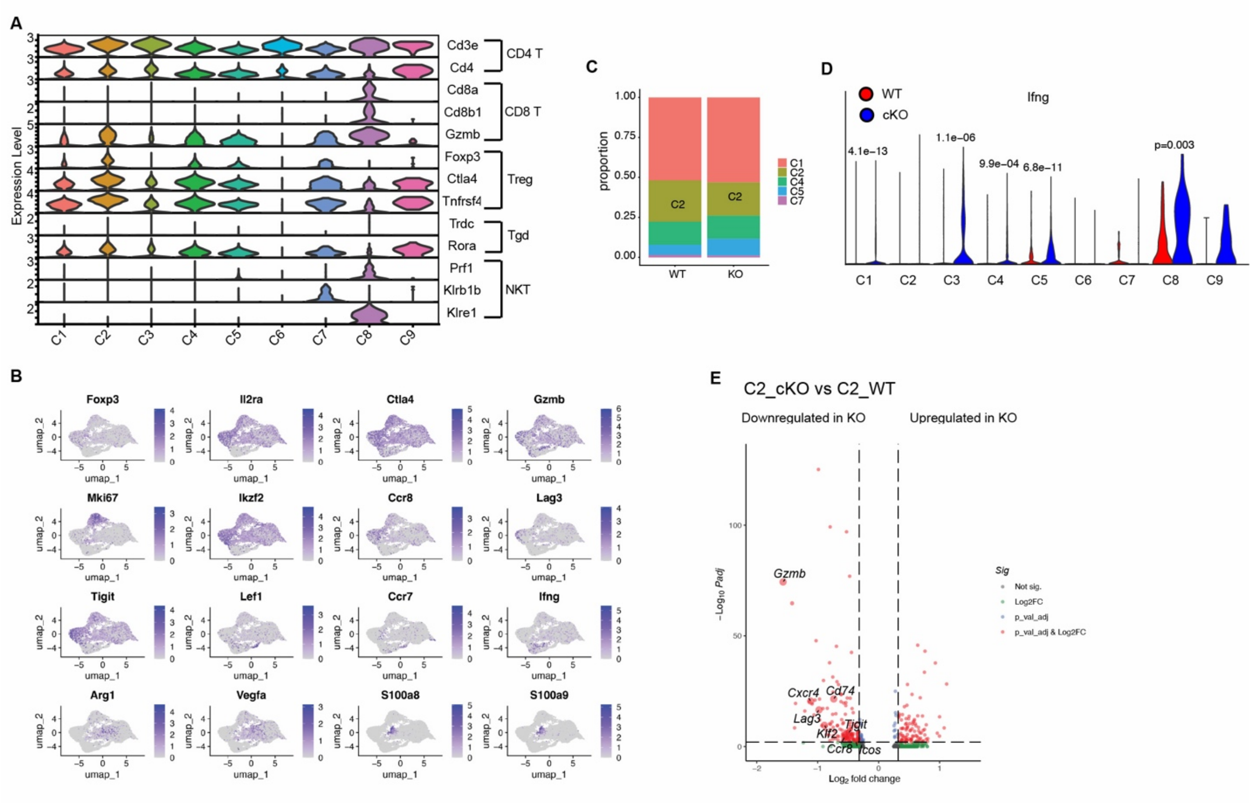

**Figure S3. scRNA-seq analysis of tumor-infiltrating CD4$^{+}$ T cells.** (A) Violin plots of canonical marker genes of each of the 9 clusters. (B) Umap plot showing gene expression within cells. (C) The proportions of Treg subsets in total Tregs. (D) Violin plot of *Ifng* expression. (E) Volcano plot comparing gene expression in C2_cKO versus WT Treg cells.

Figure S4

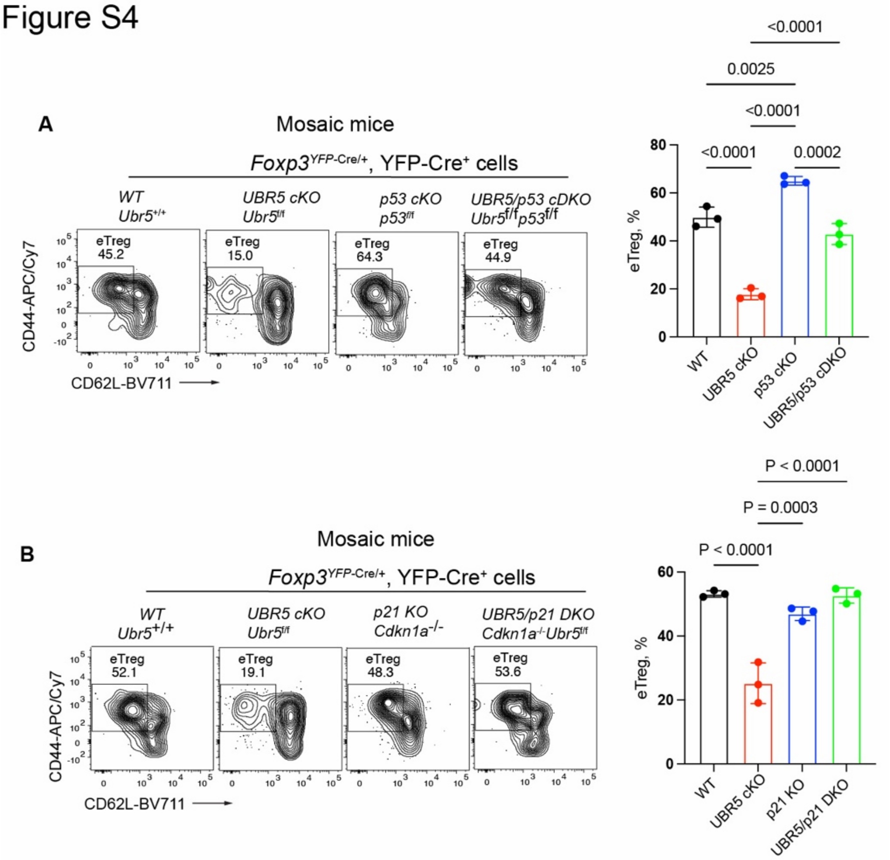


**Figure S4. UBR5 sustains Treg activation by restraining the p53–p21 checkpoint.** (A) YFP$^{+}$ Treg cells from WT (*Ubr5*$^{+/+}$), UBR5 cKO (*Ubr5*$^{f/f}$), p53 cKO (*Trp53*$^{f/f}$) and UBR5/p53 cDKO (*Ubr5*$^{f/f}$ *Trp53*$^{f/f}$) female mosaic mice were analyzed for CD44 and CD62L expression (left) and the percentages of eTregs are shown (right). (B) Tregs from WT (*Ubr5*$^{+/+}$), UBR5 cKO (*Ubr5*$^{f/f}$), p21 KO (*Cdkn1a*$^{-/-}$) and UBR5/p21 DKO (*Cdkn1a*$^{-/-}$ *Ubr5*$^{f/f}$) mosaic mice were pre-gated as YFP$^{+}$ cells and analyzed for CD44 and CD62L expression (left) and the percentages of aTregs are shown (right). N = 3 mice/group. Data are mean ± SD. Multiplicity-adjusted one-way ANOVA was used. Results are representative of two independent experiments.

Figure S5. Sorting and gating strategies

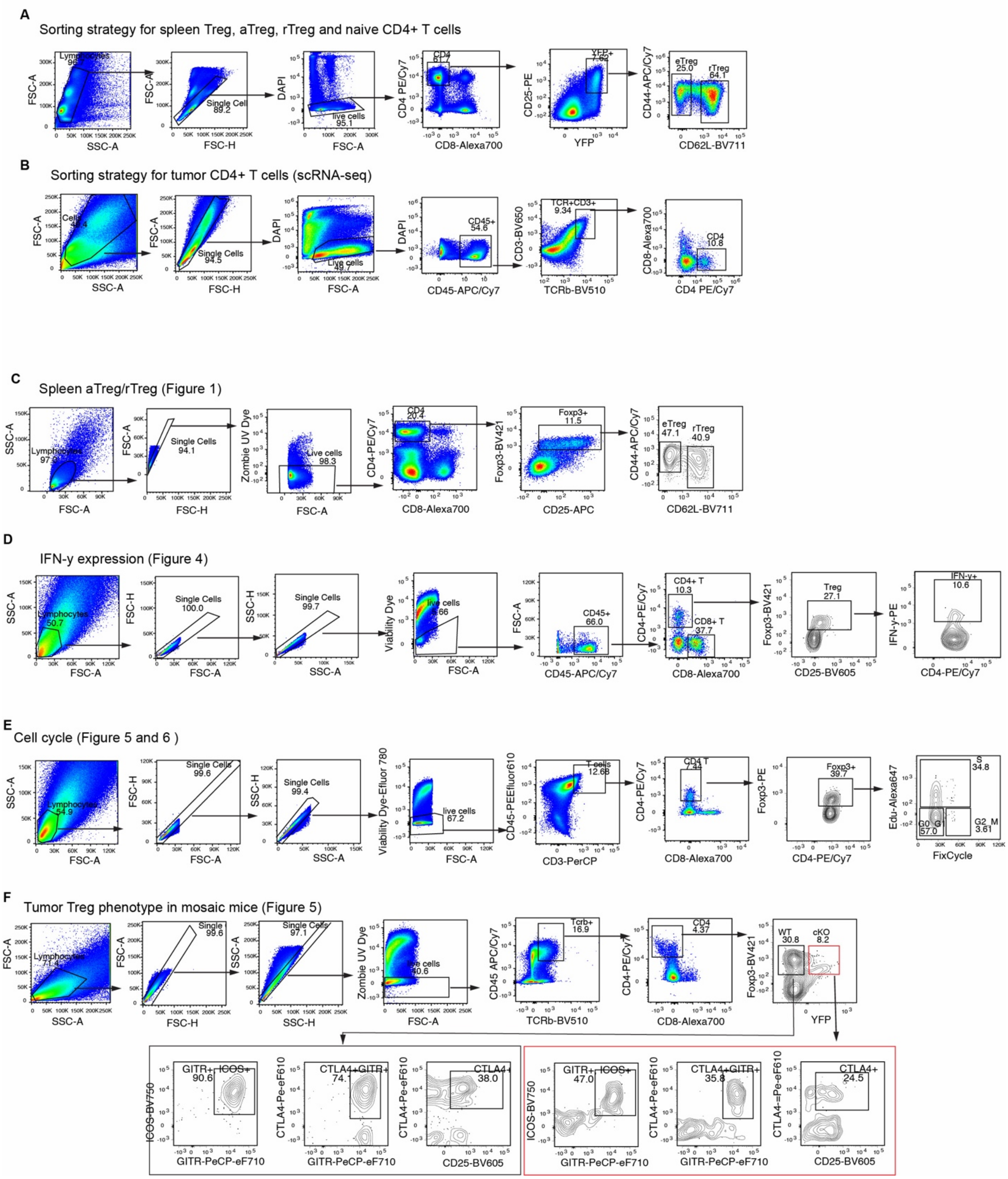


**Figure S5. Fluorescence-activated Cell Sorting (FACS) and flow cytometry gating strategies.** Sorting stratigy for eTreg, rTreg (A) and tumor-infiltrating CD4$^{+}$ T cells (B). (C) gating strategy for eTregs and rTreg in spleen. (D) gating strategy for IFN-γ expression in tumor-infiltrating Tregs.

(E) gating strategy for cell-cycle analysis in tumor Tregs. (F) gating strategy for phenotyping tumor Tregs from mosaic mice.
